# *Plagl1* is part of the mammalian retinal injury response and a critical regulator of Müller glial cell quiescence

**DOI:** 10.1101/2021.04.12.439550

**Authors:** Yacine Touahri, Luke Ajay David, Yaroslav Ilnytskyy, Edwin van Oosten, Joseph Hanna, Nobuhiko Tachibana, Lata Adnani, Jiayi Zhao, Mary Hoffman, Rajiv Dixit, Laurent Journot, Yves Sauve, Igor Kovalchuk, Isabelle Aubert, Jeffrey Biernaskie, Carol Schuurmans

## Abstract

Retinal damage triggers reactive gliosis in Müller glia across vertebrate species, but only in regenerative animals, such as teleost fish, do Müller glia initiate repair; proliferating and undergoing neurogenesis to replace lost cells. By mining scRNA-seq and bulk RNA-seq datasets, we found that *Plagl1*, a maternally imprinted gene, is dynamically regulated in reactive Müller glia post-insult, with transcript levels transiently increasing before stably declining. To study *Plagl1* retinal function, we examined *Plagl1*^+/-pat^ null mutants postnatally, revealing defects in retinal architecture, visual signal processing and a reactive gliotic phenotype. *Plagl1*^+/-pat^ Müller glia proliferate ectopically and give rise to inner retinal neurons and photoreceptors. Transcriptomic and ATAC-seq profiles revealed similarities between *Plagl1*^+/-pat^ retinas and neurodegenerative and injury models, including an upregulation of pro-gliogenic and pro-proliferative pathways, such as Notch, not observed in wild-type retinas *Plagl1* is thus an essential component of the transcriptional regulatory networks that retain mammalian Müller glia in quiescence.

## INTRODUCTION

Sight begins when photosensitive retinal neurons, the rod and cone photoreceptors, detect and convert light to electrical signals that are propagated via retinal interneurons to ganglion cells, and onto visual processing centers in the brain. Visual impairment afflicts 2.2 billion people world-wide (World Health Organization, 2019) and nearly half of these ailments are untreatable. In regenerating tissues, somatic stem cells transit from quiescent to active states to maintain tissue homeostasis and repair damage. However, most of the mammalian central nervous system (CNS), including the retina, is devoid of active stem cells and dying neurons are not replaced ^1^. Müller glia are post-mitotic, terminally differentiated somatic cells that have a host of functions under normal physiological conditions, including the provision of structural and nutrient support to other retinal cells, recycling of photopigments, and functioning as optical light fibers^2,3^. Strikingly, in teleost fish and Xenopus, Müller glia behave similar to somatic stem cells in regenerative tissues, responding to injury by first de-differentiating into proliferating progenitor cells that then acquire neurogenic competence to replace lost retinal cells ^4,5^. In contrast, very few Müller glia re-enter the cell cycle in mammals in response to injury, and fewer yet undergo neurogenesis ^6^.

A recent ground breaking study employing bulk RNA-seq, single cell (sc) RNA-seq and ATAC-seq in fish and mice in two injury models defined three Müller glial states, each upheld by distinct gene regulatory networks: resting, reactive and restoration to resting, the latter state specific to mouse ^7^. A reactive Müller glial state is triggered by injury and in both mouse and fish involves rapid and slow response genes. In both fish and mice, rapid response genes mediate TNF, MAPK, NFκB and Hippo signaling, as well as ribosome biogenesis in fish, which instead are activated as a slow response in mouse ^7^. The slow response genes in mouse involve the reversion to quiescence genes, whereas in fish, cell cycle genes are elevated, as are genes that confer neurogenic competence ^7^.

The reactive state common to mouse and fish is known as Müller glial gliosis, a stress-induced neuroprotective response characterized by cellular hypertrophy and barrier formation around the injury site, as well as the secretion of neuronal pro-survival factors, antioxidants, and pro-endothelial molecules ^6^. While moderate gliosis is transient and protective, severe gliosis is cytotoxic, and scars and remodels the retina permanently. Common features of reactive gliosis include increased expression of intermediate filament genes (e.g. GFAP, Nestin) and increased ERK signaling^6^. Other responses are stressor-specific, including reduced expression of Kcnj10/Kir4.1, a potassium channel, glutamate-ammonia ligase (Glul) (also known as glutamine synthetase), and Cdkn1b/p27Kip1, a cell cycle inhibitor ^6^.

Post-injury, mammalian Müller glia halt in a reactive state and then revert to quiescence, without proceeding to the next stage in regeneration, which is proliferation, as proliferation cues are rate-limiting ^8^. ERK signaling, which is activated downstream of growild-typeh factor stimulation, is initiated in Müller glia by insult in all species^5^, and blocking ERK prevents Müller glia proliferation in fish^9^, and the low transient proliferation seen in birds ^10^ and to a lesser extent in mammals^11,12^. Several other signals can also induce mammalian Müller glia to proliferate when ectopically activated, including Wnt ^13,14^, Notch ^13,14^, and Hedgehog^15^. Neurogenesis cues are also rate-limiting in mammals. Lineage-specifying transcription factors act in a combinatorial fashion to specify the fates of each of the seven retinal cell types ^16^. Several lineage-specifiers are upregulated post-injury in Müller glia in fish, including *Ascl1* ^17,18^, which is required for injury-induced Müller glia neurogenesis in fish ^19^, but is not induced and rate-limiting in mammalian Müller cells ^20,21^. *Ascl1* overexpression can induce neurogenesis when misexpressed in the early postnatal retina, or when combined with HDAC inhibitors, *Ascl1* can also induce adult neurogenesis, primarily promoting a bipolar cell-like fate ^22^.

Given the potential for endogenous repair based on observations in fish and frogs^4,5^, the search is on for genes that prevent mammalian Müller glia from proceeding to the proliferative and neurogenic stages of regeneration. We focused our attention on *Plagl1*, a maternally imprinted gene which, when overexpressed in the developing Xenopus ^23^ or murine ^24,25^ retina, can induce cell cycle exit. The mammalian genome contains∼150 imprinted genes, each mono-allelically expressed in a parent-of-origin-specific manner due to selective methylation of the maternal or paternal allele. Interestingly, there is growing support for the idea that even though imprinted genes encode a wide array of molecules (ncRNAs, transcription factors, signaling molecules, etc.)^26^, many have a common role in maintaining somatic stem cell quiescence, including in the lung, hematopoietic system, skeletal muscle, and brain^27-30^. Consistent with the idea that imprinted genes may have co-evolved to suit a common purpose, a meta-analyses of 85 imprinted genes revealed that they form subnetworks of co-regulated biallelically-expressed and imprinted genes^27,31^. *Plagl1* is a maternally imprinted gene encoding a zinc finger transcription factor that is part of an imprinted gene network (IGN) of 13 tightly co-regulated imprinted genes ^31^. Here we studied *Plagl1* function in the postnatal retina, revealing functional, structural, transcriptomic and epigenetic changes that culminate in an essential role for *Plagl1* in maintaining Müller glial quiescence.

## RESULTS

### *Plagl1* is expressed in Müller glial and is dynamically regulated in response to retinal damage

To determine whether *Plagl1* could function in differentiated Müller glia, we first examined transcript distribution in postnatal day (P) 7 mouse retinas, when neurogenesis and gliogenesis is complete centrally^32^. Using RNAscope *in situ* hybridization, *Plagl1* transcripts were detected in the inner nuclear layer (INL) and ganglion cell layer (GCL), overlapping with mRNA puncta for *Sox9* (Fig. 1a), a pan-glial marker expressed in Müller glia in the INL and astrocytes in the GCL^33^. To assess *Plagl1* expression at adult stages, we mined a published RNA-seq data set acquired from GFP-enriched Müller glia sorted from 2 month old *GLASTCreERT2;Sun1-sGFP* mice ^7^. We found that *Plagl1* transcript levels were enriched in GFP^+^ cells at comparable levels to other Müller glia markers (e.g., *Sox9, Vim*, Sox2, *Glul, Rlbp*), contrasting to genes marking other retinal cell types that were instead enriched in GFP-cells (e.g., *Vsx1, Vsx2, Crx, Otx2*) (Fig. 1b). *Plagl1* is thus expressed in murine Müller glia from early postnatal to adult stages.

**Figure 1.**
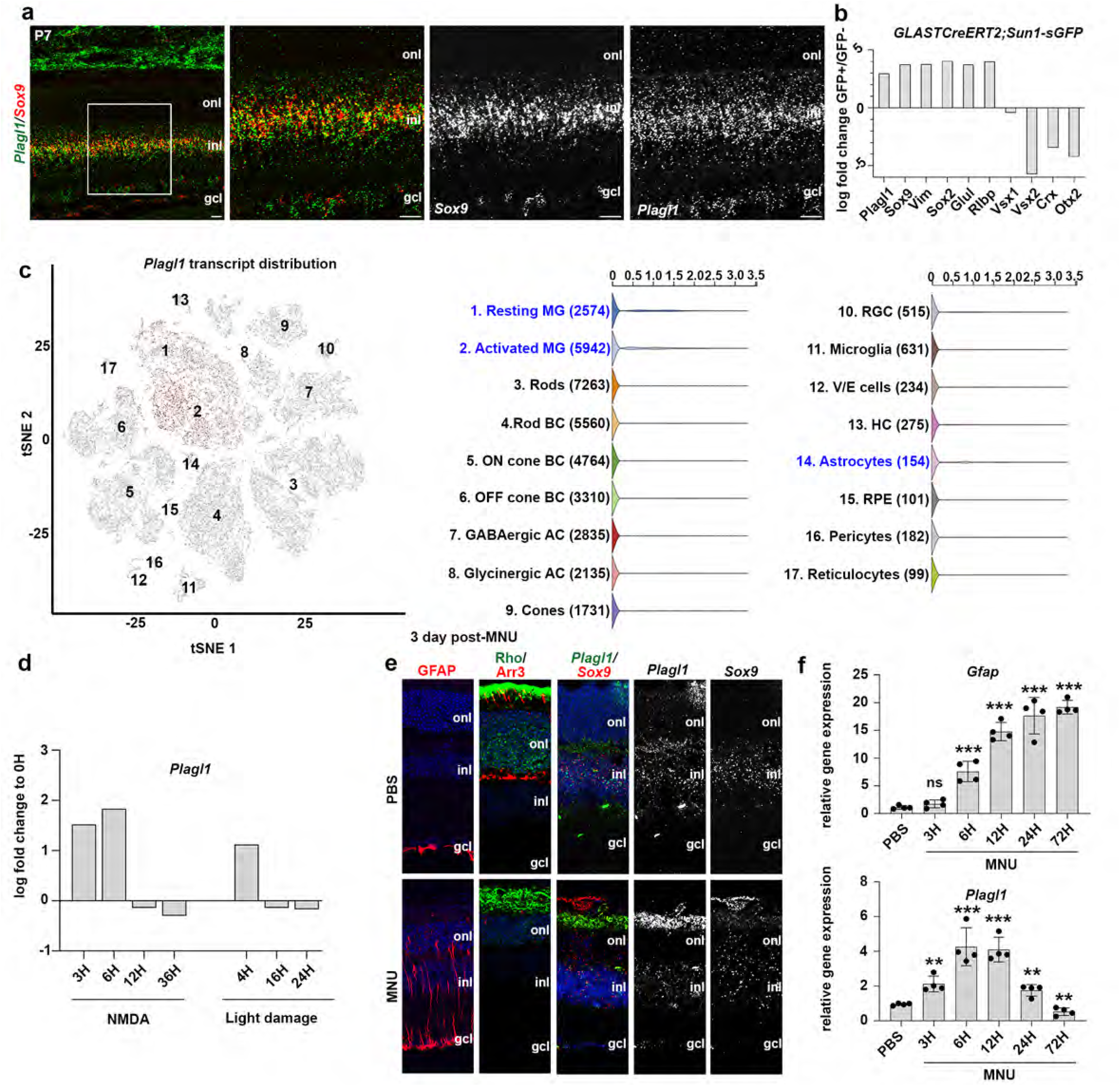
Dynamic expression of *Plagl1* in Müller glia in the healthy and injured postnatal retina. **a** RNAscope in situ hybridization of *Plagl1* (green) and *Sox9* (red) riboprobes on P7 wild-type retinal sections. Right side panels are high magnifications of boxed areas. **b** RNAseq analysis of P60 Müller glia (GFP^+^ cells sorted from *GLASTCreERT2;Sun1-sGFP* mice) extracted from the database Gene Expression Omnibus with the accession GSE135406, showing *Plagl1* expression along with Müller gli markers (*Sox9, Vim, Sox2, Glul, Rlpb*), bipolar (*Vsx1, Vsx2*) and Photoreceptor (*Crx*) markers in GFP^+^ relative to GFP-cells. **c** tSNE-dimensionality reduction to represent retinal cell clustering following NMDA treatment (data extracted from ^7^). Relative *Plagl1* expression in astrocytes= 0.108, resting Müller glia= 0.236 activated Müller glia= 0.308. **d** Temporal profiling of *Plagl1* expression in Müller glia following NMDA or light damage for 3, 6, 12, or 36 H (NMDA) or 4, 16, or 24H (light damage). RNAseq data extracted from the database Gene Expression Omnibus with the accession GSE135406. **e** Immunolabeling of GFAP, Rho, and Arr3 and RNAscope ISH for *Plagl1* and *Sox9* on adult wild-type retinal sections 3 days post PBS or MNU treatment. **f** *Gfap* and *Plagl1* expression quantified by RTqPCR in adult retinas treated with PBS or MNU for 3-, 6-, 12-, 24- and 72-hr post MNU. N=4 for all time points. Graph shows mean ± S.E.M. Unpaired Student t-test, significant differences are indicated *p < 0.05, **p < 0.01, ***p < 0.0001 *\**.gcl, ganglion cell layer; inl, inner nuclear layer; onl, outer nuclear layer. Scale bars=25μm in a, 50 μm in e.

In fish and frogs, Müller glia behave like somatic stem cells, regenerating all retinal cell types to promote repair ^4^. Conversely, mammalian Müller glia respond to insult by undergoing reactive gliosis, a stress-induced neuroprotective response characterized by cellular hypertrophy and barrier formation around the injury site ^3^. To determine how *Plagl1* expression levels change in response to injury, we first mined the comprehensive suite of scRNA-seq and bulk RNA-seq data sets performed on enriched pools of two-month-old Müller glia following NMDA and light damage (LD) injuries^7^. At the single cell level, in accordance with our RNAscope analysis, *Plagl1* was expressed in resting Müller glia and in astrocytes (Fig. 1c). However, *Plagl1* transcripts were comparatively higher in activated Müller glia, indicative of an up-regulation following retinal damage. To better characterise the dynamics of *Plagl1* expression in response to injury, we examined the distribution of *Plagl1* transcripts in bulk RNA-seq data collected from Müller glia at 3-, 6-, and 12-, and 36-hr following NMDA treatment, and at 4-, 16-, and 24-hr post LD ^7^. *Plagl1* transcript levels rapidly escalated post-injury, within 3 hr after NMDA treatment and 4 hr after LD (Fig. 1d). However, very rapidly soon after, *Plagl1* transcript levels declined below baseline, by 12-hr post-NMDA and within 16-hr post-LD (Fig. 1d). These data suggest that *Plagl1* has a dynamic response to injury, with its downregulation coinciding with the switch of Müller glia from a resting to reactive state.

To further characterise *Plagl1* expression following retinal damage, we performed our own injury, injecting 2 mo CD1 mice with N-methyl-N-nitrosourea (MNU), which rapidly induces photoreceptor degeneration ^34^. First, we confirmed the degenerative effects of MNU after 3-, 7- and 21-days post-treatment, revealing that GFAP was strongly upregulated at all stages, at the protein and transcript level, a sign that the reactive gliotic state has been triggered and is sustained (Fig. 1e, Supplementary Fig. ^1a-b^). In addition, we confirmed a partial photoreceptor loss at day 3 post-MNU, that was complete at days 7 and 21, as revealed by reduced expression of Rhodopsin and *Rcvrn*, rod markers, as well as Arr3, a cone marker (Fig. 1e, Supplementary Fig. ^1a-b^). To assess the effects of MNU on *Plagl1* expression, we first performed *Plagl1/Sox9* RNAscope, revealing an apparent decline in *Plagl1* transcript levels compared to *Sox9* (Fig. 1e, Supplementary Fig. ^1a-b^). To confirm these findings, and to further stratify the response period, we collected retinas at 3-, 6-, 12-, 24- and 72-hr and 7- and 21-days post-MNU and performed qPCR to analyse *Gfap* and *Plagl1* transcript levels (Fig. 1f, Supplementary Fig. ^1c-d^). *Plagl1* transcript levels increased within 3 hr-post MNU treatment, even before *Gfap* levels increased, and remained above baseline until 72 hr, when they began to decline, remaining below baseline up to 21-days post-injury (Fig. 1f, Supplementary Fig. ^1c-d^). Strikingly, the decline in *Plagl1* transcript levels coincides with the transitory proliferation that occurs between 48-to 60-hr post-MNU ^35^. This dynamic expression profile, especially the correlation between *Plagl1* decline and Müller glia activation and proliferation, prompted us to question whether *Plagl1* had a role in controlling Müller glia function using a loss-of-function approach.

### *Plagl1* is required to maintain retinal integrity and proper visual function

*Plagl1* is a maternally imprinted gene, resulting in mono-allelic expression from the paternal allele ^36^. To study *Plagl1* ‘loss of function’ mutants, we generated *Plagl1*^+/-pat^ mice carrying a silenced maternal allele and mutant paternal allele, these mice were previously confirmed to lack *Plagl1* expression in various tissues, including the embryonic retina ^24,27,31^. While 80% of *Plagl1*^+/-pat^ mice die in the early perinatal period^24,31^, we were able to collect surviving *Plagl1*^+/-pat^ animals at perinatal (P7), juvenile (P14, P21) and early adulthood (P60) stages. We first performed a morphological/structural assessment of the outer nuclear layer (ONL), INL and GCL using optical coherence tomography (OCT) in live P21 animals (Fig. 2a). In contrast to P21 wild-type controls, OCT images of *Plagl1*^+/-pat^ retinas revealed that layer borders were less distinct, with hypo-reflective puncta in the inner plexiform layer (ipl), indicative of layer disruption in focal areas of the retina (Fig. 2a). In transverse sections through the retina, DAPI nuclear staining confirmed these OCT observations and revealed striking retinal dysmorphologies in *Plagl1*^+/-pat^ retinas at all postnatal stages assessed (P7, P14, P21, P60), which became increasingly prevalent over time (Fig. 2b-c). Two distinct abnormalities were observed; ectopias, defined as protrusions of ONL nuclei into the subretinal space, and rosettes, defined as focal dysplasias that formed in the ONL but protruded into, and disformed, the INL (Fig. 2b, Supplementary Fig. ^2a-c^). To assess these retinal abnormalities further, we immunostained P21 wild-type and *Plagl1*^+/-pat^ retinas with molecular markers for the seven retinal cell types. The most striking disruptions were observed in the ONL, especially in the positioning of rod (Rhodospin^+^) and cone (Arr3^+^, m-opsin^+^, s-opsin^+^) outer segments, which were internalized in the central lumens of the *Plagl1*^+/-pat^ retinal rosettes (Fig. 2d, Supplementary Fig. ^3a-b^). Of the remaining cell types, amacrine (Pax6^+^, calretinin^+^, syntaxin^+^), bipolar (Ch×10^+^, PKC^+^), horizontal (calbindin^+^), and ganglion (Pax6^+^, Brn3a^+^) cell markers were grossly normal, although there were some layering disruptions in the vicinity of the *Plagl1*^+/-pat^ rosettes (Supplementary Fig. ^3c-j^). Strikingly, the morphology of Rlbp1^+^ Müller glial cell processes was also disrupted in P21 *Plagl1*^+/-pat^ retinas, especially in the rosette structures (Supplementary Fig. ^3j^).

**Figure 2.**
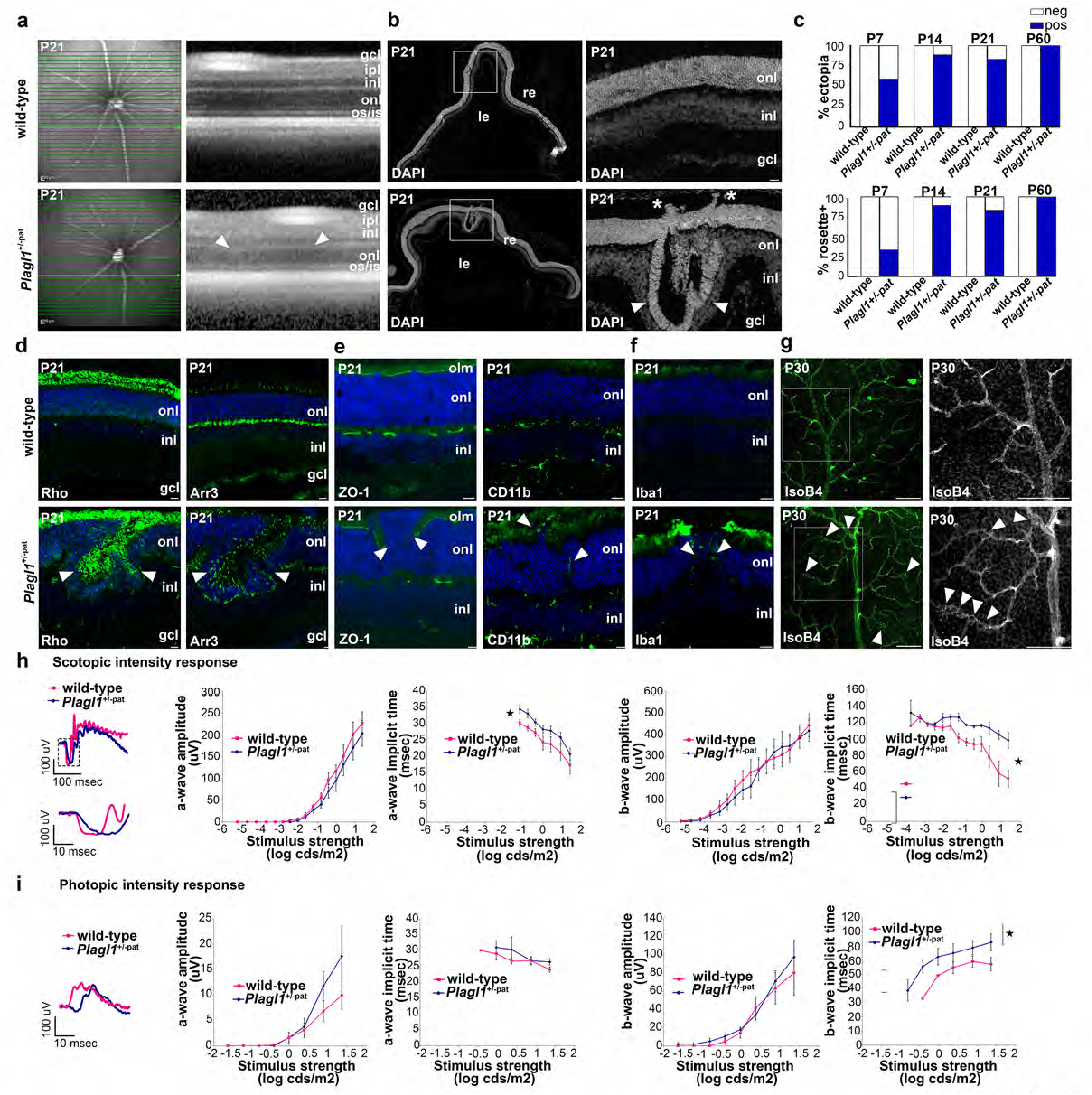
Loss of *Plagl1* function perturbs retinal architecture and function. **a** Fundus images (left) and Optical coherence tomography (OCT) derived cross-sectional images (right) of P21 wild type and *Plagl1*^+/-pat^ retinas. Green line indicates the section level. Arrow heads indicate inl disruptions. **b** DAPI staining of P21 wild-type and *Plagl1*^+/-pat^ retinas, showing retinal ectopias (asterisks) and rosettes (arrowheads). Right side panels are high magnification images of boxed areas. **c** Quantification of rosettes and ectopias in wild-type and *Plagl1*^+/-pat^ retinas at P7, P14, P21 and P60. N=5 for all stages. **d** Immunostaining for Rhodopsin and Arr3 in P21 wild-type and *Plagl1*^+/-pat^ retinal sections. Arrowheads mark ectopias and rosettes. Blue is DAPI counterstain. **e** Immunostaining for OLM marker ZO1 in P21 wild-type and *Plagl1*^+/-pat^ retinal sections. Arrowheads mark gaps in ZO-1 expression at the level of ectopias and rosettes. **f** Immunostaining for Iba1 and Cd11b with nuclear DAPI staining of P21 wild-type and *Plagl1*^+/- pat^ retinal sections. Arrowheads mark ectopias and rosettes. **g** Staining for Isolectin B4 in P30 wild-type and *Plagl1*^+m/-pat^ flatmounted retinas. Arrow heads show nodes and tortuous vessels. Right images are high magnifications of the boxed area in left images. **h, i** Full field ERG recordings of P30 wild-type and *Plagl1*^+/-pat^ mice under scotopic (**h**) and photopic (**i**) conditions. N =6 (wild-type), N =5 (*Plagl1*^+/-pat^). Plots show a- and b-wave amplitudes and implicit times. Statistics: means ± s.e.m. Student t test. *\*, p < 0*.*05; **, p < 0*.*01; ***, p <* *0*.*005*. gcl, ganglion cell layer; inl, inner nuclear layer; onl, outer nuclear layer; olm, outer limiting membrane. Scale bars: 200 μm in a (first panel), 25μm in b, d.

To characterize structural deficits in *Plagl1*^+/-pat^ Müller glia further, we examined ZO-1 staining, which marks the outer limiting membrane (OLM) of the retina, which is formed by Müller glia end-feet (Fig. 2e). ZO-1 immunostaining showed a clear disruption of OLM adherens junctions in P21 *Plagl1*^+/-pat^ retinas, especially in the vicinity of the rosettes (Fig. 2e). Strikingly, even in the absence of an injury stimulus, *Plagl1*^+/-pat^ retinas had features of an injury response. For instance, around the gaps in the OLM, Cd11b^+^ macrophages and Iba1^+^ microglia were observed to infiltrate (Fig. 2f). Finally, retinal vessels labeled with Isolectin-B4 were disorganized, forming abnormal node like structures and tortuous capillaries in P21 and P30 *Plagl1*^+/-pat^ retinas (Fig. 2g).

The disruption of *Plagl1*^+/-pat^ retinal architecture suggested that retinal activity might also be perturbed, which we assessed by recording full field electroretinograms (ERGs) in P30 juvenile *Plagl1*^+/-pat^ mice, after full visual acuity is reached around P25 ^37^ (Fig. 2h-i). Under scotopic (Fig. 2h) and photopic conditions (Fig. 2i), a and b wave amplitudes were normal in wild-type and *Plagl1*^+/-pat^ animals, indicating that the retinal circuitry remained intact in the absence of *Plagl1* function. However, implicit times for both a- and b-waves were prolonged in *Plagl1*^+/-pat^ mice under scotopic and photopic adaptation (Fig. 2h-i). Altogether, these defects are suggestive of a signal processing dysfunction rather than a global defect in retinal activity.

In summary, *Plagl1*^+/-pat^ retinas have striking morphological disruptions (ectopias, rosettes; Fig. 2j) and display signs of an injury response even in the absence of an insult. Given the prominent expression of *Plagl1* in retinal Müller glia, we focused on how these glial cells are affected by the loss of *Plagl1* for the remainder of this study.

### *Plagl1* mutant Müller glia undergo insult-independent reactive gliosis

The morphological defects observed in postnatal *Plagl1*^+/-pat^ retinas, were strikingly reminiscent of other mutations that result in a loss or dysfunction of mature Müller glia^38-40^, in keeping with the known structural role of these glial cells ^3,41^. To determine whether *Plagl1* is required for Müller glia homeostasis, we searched for signs of reactive gliosis, an ‘activation’ state that normally occurs in response to insult or pathology that is associated with increased expression of intermediate filaments (GFAP, Vimentin) and a decline in Glul activity. In P14 wild-type controls, Vim labeled Müller glia cell bodies, processes, and endfeet, especially those that extend to the GCL (Fig. 3a). Vim immunoreactivity increased in *Plagl1*^+/-pat^ retinas, with outer processes and endfeet now clearly labeled, especially in the vicinity of the rosette-like structures (Fig. 3a). Similarly, GFAP expression, which was only expressed in astrocytes lining the GCL in wild-type retinas, was up-regulated in *Plagl1*^+/-pat^ retinas, beginning at P14, but more strikingly obvious at P21 and P45, especially in the vicinity of the retinal rosettes (Fig. 3b). We also confirmed the reactive state of *Plagl1*^+/-pat^ Müller glia by examining ERK phosphorylation on T202/Y204 (designated pERK), which is normally triggered by growild-typeh factor signaling in reactive Müller glia. pERK protein was detected in Müller glial processes in wild-type retinas, but there was a striking upregulation in expression in cell bodies, including ectopic glia in the ONL, and in glial processes, in P14 *Plagl1*^+/-pat^ retinas, which we confirmed by Western blotting (Fig. 3c,e). Finally, reactive gliosis is also characterized by a reduction of Glul expression, which we also observed in *Plagl1*^+/-pat^ retinas, with gaps in labeled Müller glia in the INL, a reduction that was verified by Western blotting (Fig. 3d,f).

**Figure 3.**
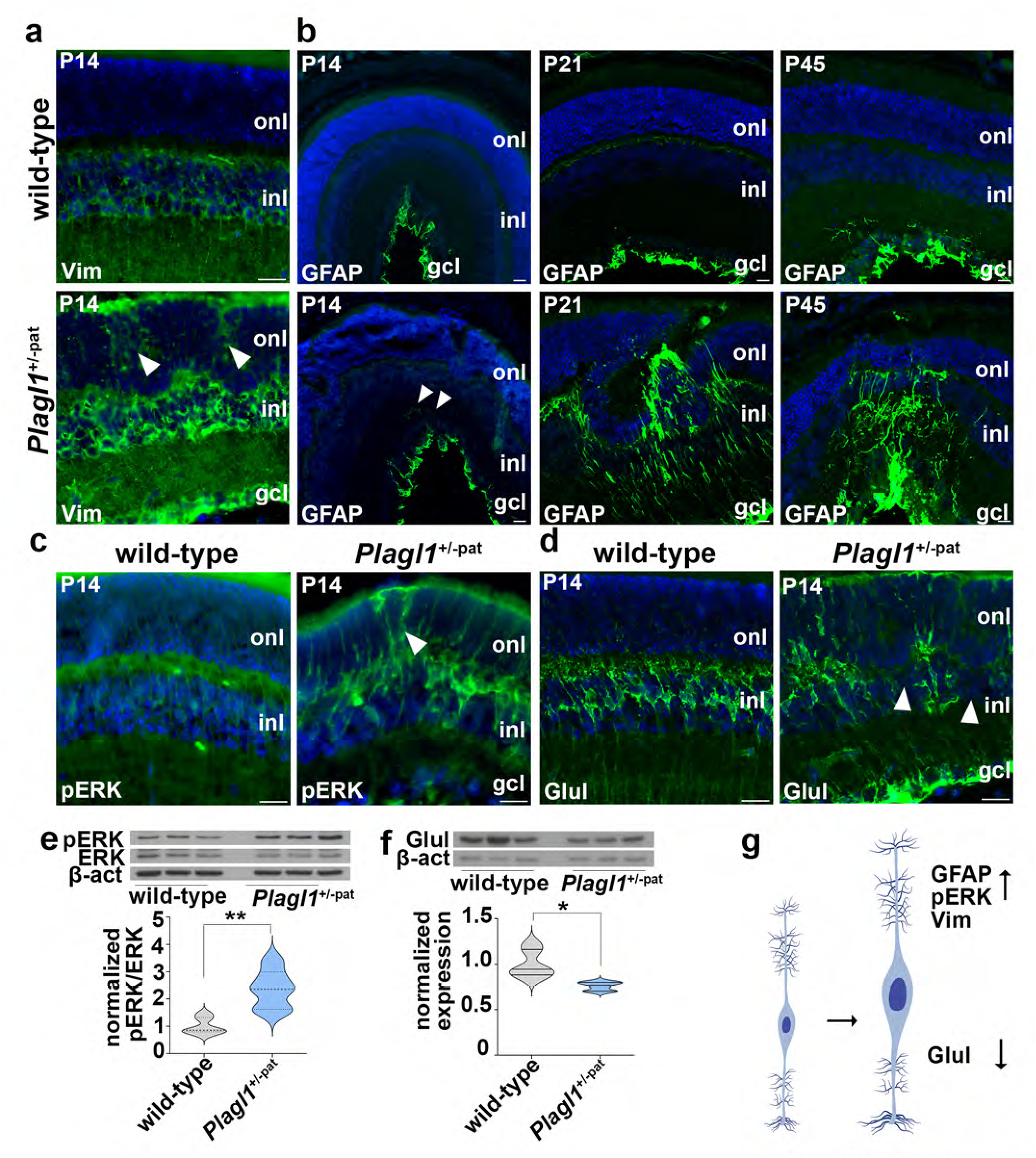
Loss of *Plagl1* triggers spontaneous reactive gliosis. **a** Immunostaining for vimentin (Vim) in P14 wild-type and *Plagl1*^+/-pat^ retinal sections. Arrow - heads point to ectopic Vim upregulation in *Plagl1*^+/-pat.^ **b** Immunostaining for GFAP in P14, P21 and P45 wild-type and *Plagl1*^+/-pat^ retinal sections. Arrowheads point to ectopic GFAP expression in *Plagl1*^+/-pat^ retinas. **c** Immunostaining for pERK in P14 wild-type and *Plagl1*^+/-pat^ retinal sections. Arrowheads point to pERK upregulation in *Plagl1*^+/-pat^ retinas. **d** Immunostaining for Glul staining in P14 wild-type and *Plagl1*^+/-pat^ retinal sections. Arrowheads point to Glul gaps in *Plagl1*^+/-pat^ retinas. **e** Western blotting for pERK in P14 wild-type and *Plagl1*^+/-pat^ retinas showing increase in pERK protein levels. N=6 for both wild-type and *Plagl1*^+/-pat^ retinas. **f** Western blotting for Glul in P14 wild-type and *Plagl1*^+/-pat^ retinas showing decrease of Glul protein levels. N=6 for both wild-type and *Plagl1*^+/-pat^ retinas. **g** Schematic of the molecular changes associated with reactive gliosis observed in *Plagl1*^+/-pat^retinas. Statistics: means ± s.e.m. Student t-test. *\*, p < 0*.*05; **, p < 0*.*01; ***, p < 0*.*005*. gcl, ganglion cell layer; inl, inner nuclear layer; onl, outer nuclear layer. Scale bars: 25μm.

Taken together, these analyses suggest that *Plagl1*^+/-pat^ Müller glia display several hallmark features of reactive gliosis (Fig. 3g), even in the absence of injury.

### *Plagl1* mutant Müller glia undergo interkinetic nuclear migration and proliferate ectopically

Another hallmark feature of Müller glia activation in regenerative species is that glial nuclei undergo interkinetic nuclear migration (INM), moving into the ONL where they then divide to initiate the proliferative phase of the repair process ^42^. To assess whether Müller glial undergo INM in *Plagl1*^+/-pat^ retinas, we used Sox9, which marks Müller glia nuclei^43^. While in P21 wild-type retinas, Sox9^+^ cells formed essentially a monolayer in the INL, in *Plagl1*^+/-pat^ retinas, Sox9^+^ glial nuclei were detected in the INL and in the ONL, especially in the vicinity of the retinal rosettes and ectopia (Fig. 4a,b). Moreover, there was an increase in the total number of Sox9^+^ nuclei in *Plagl1*^+/-pat^ retinas, at least in the ONL in the vicinity of the rosettes (Fig. 4b), where an injury-like response has been triggered (see Figs. 2, 3).

**Figure 4.**
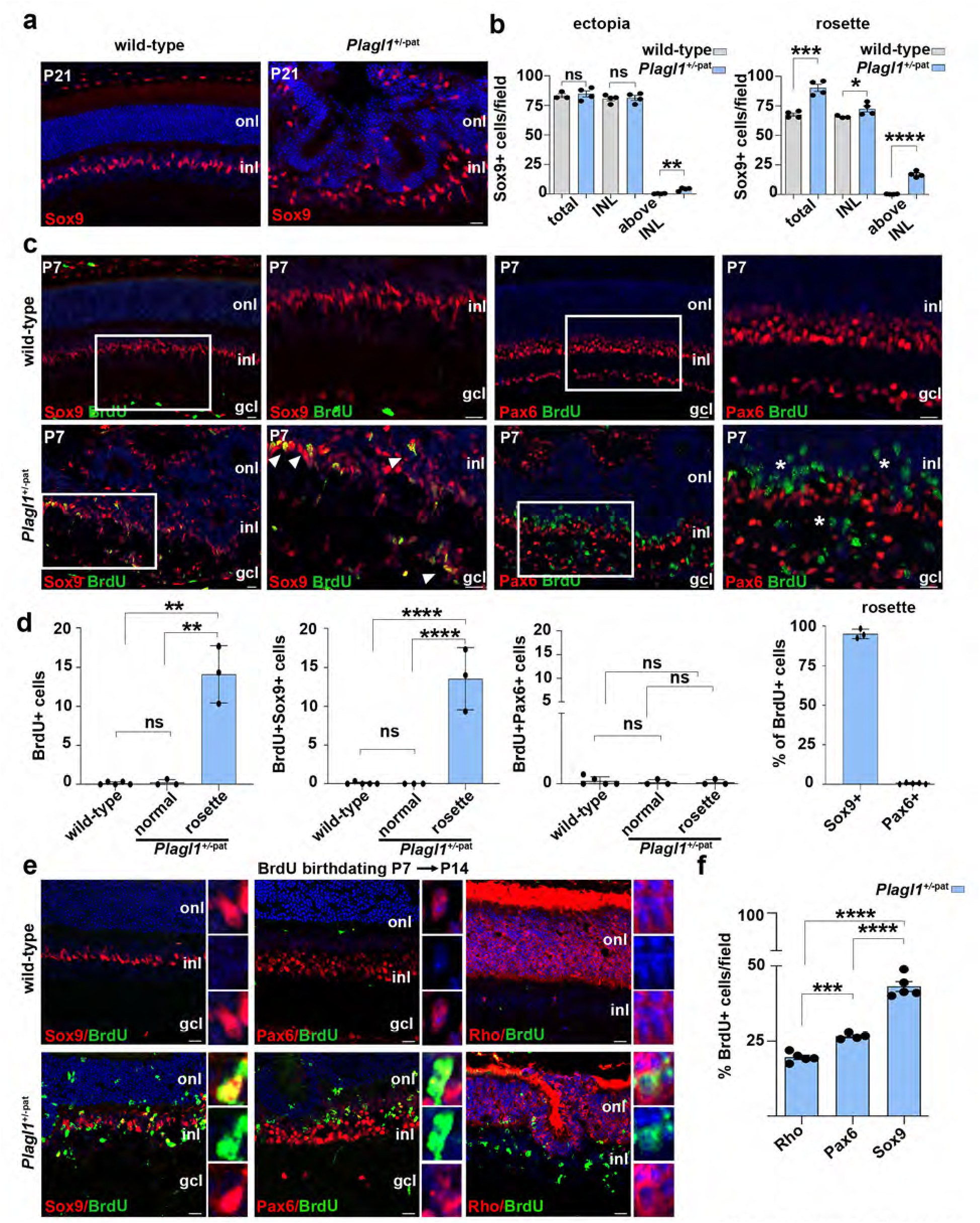
*Plagl1*^m/-p^ Müller glia proliferate ectopically. **a** Immunostaining for Sox9 in P21 wild-type and *Plagl1*^+m/-pat^ retinal sections showing increase and mis-localisation of Sox9^+^ cells in *Plagl1*^+m/-p^ retinas. Blue is DAPI counterstain. **b** Quantification of total, INL, and above INL located Sox9^+^ cells number in P21 wild-type and *Plagl1*^+m/-pat^ ectopia and rosettes-containing retinal sections. N=4 for wild-type and *Plagl1*^+/-pat^ retinas. **c** Immunostaining for Sox9, Pax6, and BrdU on P7 wild-type and *Plagl1*^+m/-pat^ retinal sections injected with BrdU for 30 min prior to harvest. Blue is DAPI counterstain. Right images are high magnifications of the boxed area in left images. Arrowheads point to BrdU positive Sox9 cells and asterisks point to BrdU negative for Pax6 in *Plagl1*^+/-pat^ retinas. **d** Quantification of BrdU^+^ cells, and the percentage of BrdU^+^ cells that are BrdU^+^Sox9^+^ and BrdU^+^Pax6^+^ in P7 wild-type and *Plagl1*^*+m/-pat*^ normal and rosettes-containing retinal sections injected with BrdU 30 min prior to dissection at P7. wild-type: N=5, *Plagl1*^+/-pat^ N=3. **e** BrdU birthdating (P7 BrdU to P14 dissection) *in vivo*, followed by BrdU co-immunostaining with Sox9, Pax6, and Rhodopsin of P14 wild-type and *Plagl1*^+m/-pat^ retinal sections. Blue is DAPI counterstain. **f** Quantification of BrdU birthdating (P7 BrdU to P14 dissection), showing the percentage of BrdU^+^ cells that are BrdU^+^Rho^+^, BrdU^+^Pax6^+^ and BrdU^+^Sox9^+^ in P14 wild-type and *Plagl1*^*+m/-pat*^ normal and rosettes-containing retinal sections *in vivo*. wild-type: N=5, *Plagl1*^+/-pat^ N=5. Statistics: mean ± S.E.M, Mann–Whitney U-test. significant differences: *p<0.05, **p<0.01, ***p <0.0001. gcl, ganglion cell layer; inl, inner nuclear layer; onl, outer nuclear layer. Scale bars: 25μm.

Terminally differentiated Müller glia are normally quiescent. Given the increase in Sox9^+^ nuclei in *Plagl1*^+/-pat^ retinas, we questioned whether these glial cells had aberrantly entered the cell cycle. To examine the proliferative response, we analyzed BrdU incorporation into dividing S-phase nuclei in P7 retinas, when differentiation in the central retina is complete, at P14, when differentiation is complete throughout the retina, and at P21 ^32^. BrdU^+^ cells were detected at P7 in the central retinas of *Plagl1*^+/-pat^ animals and not their littermate controls, but only in the dysmorphic areas containing clear rosette-like structures (Fig. 4c,d). To identify these proliferating cells, we co-stained with BrdU and Sox9, a Müller glia marker^43^. We found that 95% of the BrdU^+^ cells were Sox9^+^ in P7 *Plagl1*^+/-pat^ retinas, suggesting that the majority of dividing cells are glia (Fig. 4c,d). However, because Sox9 is initially expressed in retinal progenitor cells (RPC) and only later becomes Müller glia restricted ^43^, to confirm that these dividing cells were not a retained pool of proliferating RPCs, we also co-stained with the RPC marker Pax6^44^. Only 0.4% of the BrdU^+^ cells detected in *Plagl1*^+/-pat^ retinas expressed Pax6 in the rosettes (Fig. 4c,d). Thus, the vast majority of proliferating cells in the P7 *Plagl1*^+/-pat^ retina express Sox9, a Müller glia marker, and not Pax6, an RPC marker.

### *Plagl1* mutant retinas give rise to new neurons after the normal neurogenic period is over

In response to injury, Müller glia in all species first undergo reactive gliosis, but only in regenerative species, such as teleost fish and frogs, do these cells re-enter the cell cycle and then undergo neurogenesis ^7^. To determine whether proliferating cells in P7 *Plagl1*^+/-pat^ retinas, which are mainly glia, have also acquired neurogenic competence, we performed a birthdating assay. Proliferating cells were labeled with BrdU at P7, and the fate of these cells was then analyzed at P14 by staining with cell type-specific markers, including Sox9 for Müller glia, Pax6 for amacrine, horizontal and ganglion cells, and Rhodopsin for rods (Fig. 4e). As a thymidine analog, BrdU is incorporated into dividing nuclei in S-phase, and while diluted out of rapidly dividing cells, it permanently labels their non-dividing progeny. Consistent with the 30 min BrdU assays conducted to assess proliferation (Fig. 4c), we only observed BrdU^+^ cells in *Plagl1*^+/-pat^ and not in wild-type controls in the central retina (Fig. 4e). Strikingly, only 45% of BrdU^+^ cells stained for the glial marker, Sox9, suggesting that the proliferating cells in the central retina had given rise to other retinal cells (Fig. 4f). Indeed, 23% of the BrdU^+^ cells were Pax6^+^, while 19% localized to the ONL and expressed Rhodopsin, a rod marker (Fig. 4f).

In aggregate, these data suggest that *Plagl1* is an essential negative regulator of Müller glia activation, including the proliferative response and neurogenic potential of these cells. As *Plagl1* encodes a transcriptional regulator, we next sought to identify downstream transcriptional changes in *Plagl1*^+/-pat^ retinas that may explain these changes.

### *Plagl1* regulates the expression of a large network of retinal genes

To decipher the molecular mechanisms underlying the requirement for *Plagl1* to maintain Müller glia quiescence, we performed bulk RNA-seq on P7 wild-type control and *Plagl1*^+/-pat^ retinas, at a stage when these glial cells were proliferating ectopically in *Plagl1*^+/-pat^ retinas. Global transcriptomic analysis identified 2552 differentially expressed genes (DEGs) in *Plagl1*^+/-pat^ retinas, including up-(1410) and down-(1142) regulated genes, as expected given that *Plagl1* can act as a transcriptional co-/activator or co-/repressor^45-48^ (Fig. 5a). To confirm that P7 *Plagl1*^+/-pat^ animals were true null mutants, with a silenced maternal allele, we examined *Plagl1* expression levels. *Plagl1* was among the most downregulated genes, but transcripts were only reduced by 50%. However, a survey of read coverage revealed that *Plagl1* exons 5 and 6, covering deleted regions in the mutant allele, were missing in P7 *Plagl1*^+/-pat^ retinas (Fig. 5b), confirming the lack of maternal transcription, as previously shown in *Plagl1*^+/-pat^ mouse embryonic fibroblasts (MEFs)^49^.

**Figure 5.**
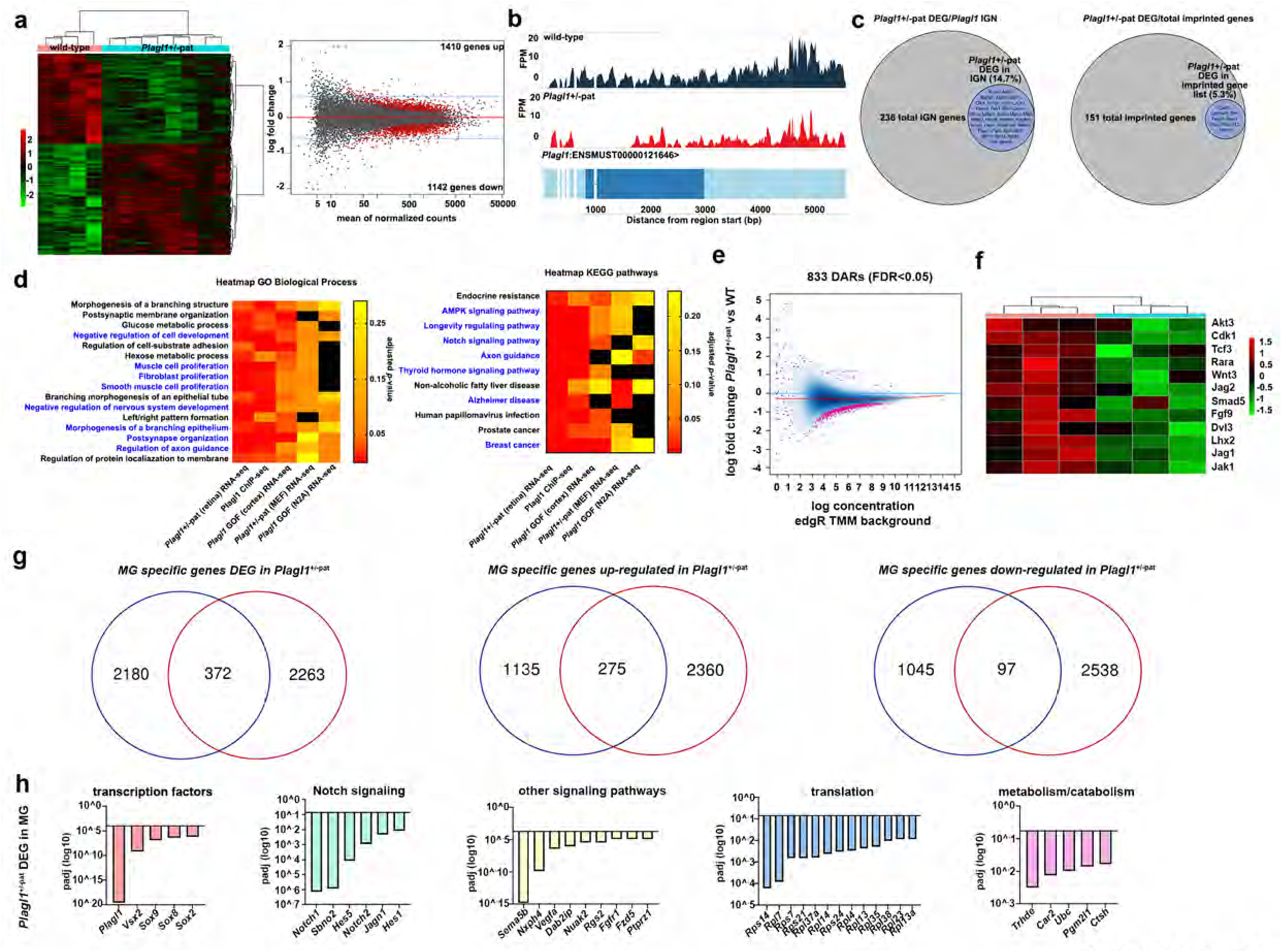
Transcriptomic and epigenomic profiling of *Plagl1*^+m/-p^ retinas. **a** Heatmap and MAplot showing genes differentially expressed in P7 *Plagl1*^+m/-pat^ retina. **b** RNaseq read coverage of *Plagl1* genomic locus in wild-type and *Plagl1*^+m/-pat^ showing absence of coverage of exon 2 in *Plagl1*^+m/-pat^ indicating that all the sequenced transcripts are of paternal origin. **c** Venn diagram identifying the proportion of IGN and all imprinted genes within *Plagl1*^+m/-pat^DEGs. **d** GO biological processes and KEGG pathways analysis comparing *Plagl1*^+m/-pat^ retinal transcriptome to *Plagl1* CHIP-seq, *Plagl1* GOF in the cortex, *Plagl1*^+m/-pat^ MEF transcriptome, and *Plagl1* GOF in N2A cell data sets. **e** ATACseq MAplot showing differentially accessible chromatin regions in P7 *Plagl1*^+m/-pat^ vs wild-type retinas. **f** Chromatin accessibility heatmap of genes corresponding to DARs that are differentially expressed in *Plagl1*^+m/-pat^ retinas. **g** Venn diagram identifying Müller glia specific genes that are differentially expressed in *Plagl1*^+m/-pat^, followed by Müller glia specific genes that are up and down-regulated in *Plagl1*^+m/- pat^ retinas. **h** Characterisation of Müller glia specific genes that are differentially expressed in *Plagl1*^+m/-pat^ retinas. Enriched GO categories include: transcription factors, Notch signalling, other signalling pathways, translation, and metabolism/catabolism.

*Plagl1* is part of an imprinted gene network (IGN) of biallelically and mono-allelically expressed genes that are tightly co-regulated, the functions of which are poorly understood in the retina in general and in Müller glia in particular^31^. Strikingly, 35/238 IGN genes (15%) were differentially expressed in *Plagl1*^+/-pat^ retinas (Fig. 5c). Furthermore, a scan of other known imprinted genes (151) identified an additional 6 that were deregulated in *Plagl1*^+/-pat^ (*Ube3a, Casd1, Commd1, Tbc1d12, Rian, Peg10*) (Fig. 5c), confirming the role of *Plagl1* as a critical regulator of the IGN and highlighting its essential function as a co-regulator of imprinted genes in the retina.

To further classify *Plagl1-*regulated pathways in the retina, we examined gene ontology (GO) term enrichment in the DEG dataset (Supplementary Fig. ^4a^). Included amongst the top 30 GO terms in Molecular Function were several terms associated with transcriptional regulation (e.g., “*transcription coregulator activity*”, “*DNA-binding transcription factor binding*”), while the top 30 Biological Processes enriched GO terms were associated with chromatin modification (e.g. “*covalent chromatin modification*”, “*histone modifications*”), neuronal differentiation (e.g. “*axogenesis*”, “*positive regulation of neuron differentiation*”), and neural development (“*stem cell maintenance*”, “*maintenance of cell number*”) (Supplementary Fig. ^4a^). Similar GO terms were enriched in Cellular Component analyses, including several terms associated with neuronal differentiation (e.g., “*neuron-to-neuron synapse*”, “*post-synaptic density*”) (Supplementary Fig. ^4a^). Finally, we also examined Kyoto Encyclopedia of Genes and Genomes (KEGG) pathways, revealing an enrichment of genes associated with neurodegeneration (e.g., “*Alzheimer’s disease*”, “*Huntington’s disease*”, “*Parkinson’s disease*”), cancer (e.g. “*breast cancer*”, “*prostate cancer*”, “*pancreatic cancer*”) (Supplementary Fig. ^4a^).

In summary, the analysis of *Plagl1*^+/-pat^ DEG in the P7 retina suggested that several cellular and molecular processes could have gone awry, including those involved in chromatin and transcriptional regulation, neural development and neuronal differentiation, and neurodegeneration, and cancer.

### *Plagl1* regulates a common subset of genes and pathways in different cell types

Consistent with our finding that *Plagl1* maintains Müller glia quiescence, human *PLAGL1* is a tumour suppressor gene, and induces cell cycle exit when misexpressed in cell lines ^50-53^ or in the developing nervous system ^24,54^. To narrow down genes/pathways potentially responsible for the ectopic Müller glia proliferation observed in P7 *Plagl1*^+/-pat^ retinas, we compared retinal DEGs to those identified in previous Plagl1 ChIP-seq and *Plagl1* loss- and gain-of-function (GOF) RNA-seq studies performed in other cell types and tissues (Fig. 5d). Our goal was to identify genes/pathways commonly regulated by *Plagl1* in different cellular contexts. Plagl1 ChIP-seq using *Plagl1-*overexpressing Neuro-2a cells identified 3859 distinct gene loci bound by Plagl1, in vast excess of the 351 *Plagl1*-regulated genes in these cells, suggesting that Plagl1-binding is not always associated with gene transcription^49^. Nevertheless, we found that 557/3859 (14.4%) of Plagl1-bound gene loci in Neuro2a cells were amongst the DEGs in P7 *Plagl1*^+/-pat^ retinas, while 34/351 (9.7%) of the genes induced by *Plagl1* overexpression in Neuro2a cells were also de-regulated in *Plagl1*^+/-pat^ retinas (Fig. 5d). An even larger proportion of DEGs in P7 *Plagl1*^+/-pat^ retinas were shared with those that were deregulated when *Plagl1* was overexpressed in embryonic cortical progenitors (232/1400 genes = 16.6%) ^55^ (Fig. 5d). Finally, a comparison to RNA-seq data of *Plagl1*^+/-pat^ mouse embryonic fibroblasts (MEFs) ^49^ identified 36/238 genes (15.1%) as commonly deregulated in the P7 *Plagl1*^+/-pat^ retina (Fig. 5d).

We next assessed GO term and KEGG pathway enrichment in the commonly deregulated genes in each of these datasets (Fig. 5d; Supplementary Fig. ^4b^). Of the GO biological process terms, proliferation was a top de-regulated term amongst retinal DEGs and the Plagl1 ChIP-seq data, while several neuronal differentiation terms were also shared with *Plagl1*^+/-pat^ retinas. One neuronal differentiation GO term was also significantly enriched amongst the DEG in *Plagl1*^+/-pat^ mutant retinas, Plagl1 ChIP-seq data and *Plagl1* GOF in cortex. Finally, KEGG pathway analysis identified Notch signaling and various cancer terms as the only shared pathways between *Plagl1*^+/- pat^ mutant retinas, Plagl1 ChIP-seq and *Plagl1* GOF (padj<0.05).

Taken together, these comparative studies suggest that *Plagl1* controls common gene sets across cell types, which could be involved in retinal homeostasis and tissue morphogenesis, as well as proliferation and neuronal differentiation, processes that we investigated further below in *Plagl1*^+/-pat^ retinas.

### *Plagl1* globally reduces chromatin accessibility without altering the expression of critical glial genes

“Covalent chromatin modification” and “histone modification” were the two top GO biological processes associated with DEGs in P7 *Plagl1*^+/-pat^ retinas (Supplementary Fig. ^4a^). To test whether the transcriptomic changes observed in *Plagl1*^+/-pat^ are due to epigenetic regulation, we used ATAC-seq to identify nucleosome-sparse (open) genomic DNA^56^. ATAC-seq was performed on P7 control and *Plagl1*^+/-pat^ retinas, which identified 833 differentially accessible regions (DARs). Strikingly, the vast majority of the *Plagl1*^+/-pat^ DARs (814/833) displayed decreased accessibility in *Plagl1*^+/-pat^ retinas, while 19 had increased accessibility (Fig. 5e). We compared DARs and DEGs in *Plagl1*^+/-pat^ retinas by combining ATACseq and RNAseq data. We limited the DARs list to 311 peaks located within 5000 bp from the nearest transcription start site (TSS) and compared to the RNAseq data for the associated genes. As expected, the majority of DARs (308/311) were associated with the decrease of accessibility in *Plagl1*^+/-pat^. While no increased DARs (0/3) overlapped with either up- or down-regulated genes, out of 308 decreased accessibility DARs, 15 overlapped with down-regulated genes and 36 with up-regulated genes in *Plagl1*^+/-pat^ (Fig. 5f). Interestingly, several of the upregulated genes were identified as genes involved in various aspects of Müller glia homeostatic functions and/or the injury response (Fig. 5f).

Taken together, these comparative studies suggest that *Plagl1* controls, at the transcriptional level, gene sets involved in retinal homeostasis, which we investigated further below in *Plagl1*^+/-pat^ retinas.

### Additional signs of a retinal injury occur in *Plagl1*^+/-pat^ retinas

Müller cell gliosis occurs in response to retinal injury or disease, and in chronic conditions, contributes to a neurodegenerative process^57^. Activation of Müller glia in *Plagl1*^+/-pat^ mutants suggested that there may be additional shared features with other retinal degeneration models. We thus compared the *Plagl1*^+/-pat^ transcriptome to neurodegenerative models (Supplementary Fig. ^4c-e^). We first examined Norrin (*Ndp*) knock-outs (KOs), which are a model of retinal hypo-vascularization and chronic hypoxia due to a disruption of the blood-retinal-barrier (BRB), and which also display reactive gliosis ^58^. Of the 92 Müller glia enriched genes in *Ndp* KOs, 19 (20.7%) genes were among the DEGs in P7 *Plagl1*^+/-pat^ retinas, including Vegf, a critical regulator of angiogenesis. In *rd10* mutants, which undergo a progressive loss of rod photoreceptors, of the 20 Müller glia-enriched genes, 4 (20%) were also deregulated in P7 *Plagl1*^+/-pat^ retinas (Supplementary Fig. ^4d^). Included was *Vim*, a marker of reactive gliosis, as well as *Sox9*, a glial-specific retinal gene that is upregulated in injury^8^. Finally, in *Rds/Prph2* mutant mice, treatment with CNTF induced GFAP expression and reactive gliosis, and transcriptomic analyses identified 4 Müller glia-enriched genes ^59^, including *Vsx2*, a retinal progenitor cell (RPC) gene upregulated in injury, which was amongst the in P7 *Plagl1*^+/-pat^ retinal DEGs (Supplementary Fig. ^5e^).

The transcriptomic resemblance of *Plagl1*^+/-pat^ retinas to neuronal degeneration models and injury prompted a more detailed comparison to the recent comprehensive analysis of the Müller glia response to injury ^7^.

### *Plagl1* controls a subset of Müller glia specific genes

The Müller glia proliferation phenotype in addition to the expression profile suggest a prominent role of *Plagl1* in maintaining Müller glia in quiescence. To better understand *Plagl1* function, we analysed *Plagl1*^+/-pat^ DEGs that were expressed in Müller glia sorted from *GLASTCreERT2;Sun1-sGFP* mice^7^. In this large dataset, of 2263 genes expressed in Müller glia, 372 were among the P7 *Plagl1*^+/-pat^ retinal DEGs (Fig. 5g). Among these 372 DEGs, 275 were up-regulated while 97 were down-regulated (Fig. 5g). Included were glial determinants of the Nfi (*Nfib, Nfix*) ^60^ and Sox (*Sox2, Sox4, Sox6 Sox8, Sox9*) ^61^ families, the hippo pathway effectors *Tead1* and *Tead2*, Notch pathway genes *(Notch1, Notch2, Hes1, Hes5, Snbo2*), other signalling pathways (*Sema5b, Rgs2, Fgfr1, Fzd5*), ribosomal genes involved in translation (*Rps19, Rps5, Rps4x, Rpl23, Rpl14, Rpl7l1, Rpl35, Rpl29*) and metabolism/catabolism related genes (*Trhde, Car2, Ubc, Ctsh*) (Fig. 5h).

To further characterize the state of *Plagl1*^+/-pat^ Müller glia, we compared DEGs to the transcriptomes of NMDA and LD injured retinas. Comparing DEGs in the 3 conditions showed that of the 2552 genes differentially expressed in *Plagl1*^+/-pat^ retinas, 637 are injury responding genes that are also DEGs in both NMDA and LD treatments (Fig. 6a). Next, we compared the behaviour of these differentially expressed genes in *Plagl1*^+/-pat^ and injured Müller glia at different time points. We identified: (1) genes that were altered in the same direction in *Plagl1*^+/-pat^ and injured Müller glia (up-or down-regulated), (2) genes that were differentially expressed in injury and not in *Plagl1*^+/-pat^ retinas (or vice versa), and (3) genes that were up-regulated in *Plagl1*^+/-pat^retinas and down-regulated in injury (or vice versa) (Fig. 6b, Supplementary Fig. ^5a^). Interestingly, we found that several Notch pathway genes were upregulated in P7 *Plagl1*^+/-pat^ mutant retinas (Fig. 6c, Supplementary Fig. ^5b^,^c^) whereas they were downregulated in NMDA and LD injured Müller glia (Fig. 6d). Included were Notch downstream effectors, *Hes1* and *Hes5*, the receptors *Notch1* and *Notch2*, and the ligand *Dll1*, while *Serr* and *Jag2* ligands were only up-regulated in *Plagl1*^+/-pat^ (Fig. 6c,d, Supplementary Fig. ^5b^,^c^).

**Figure 6.**
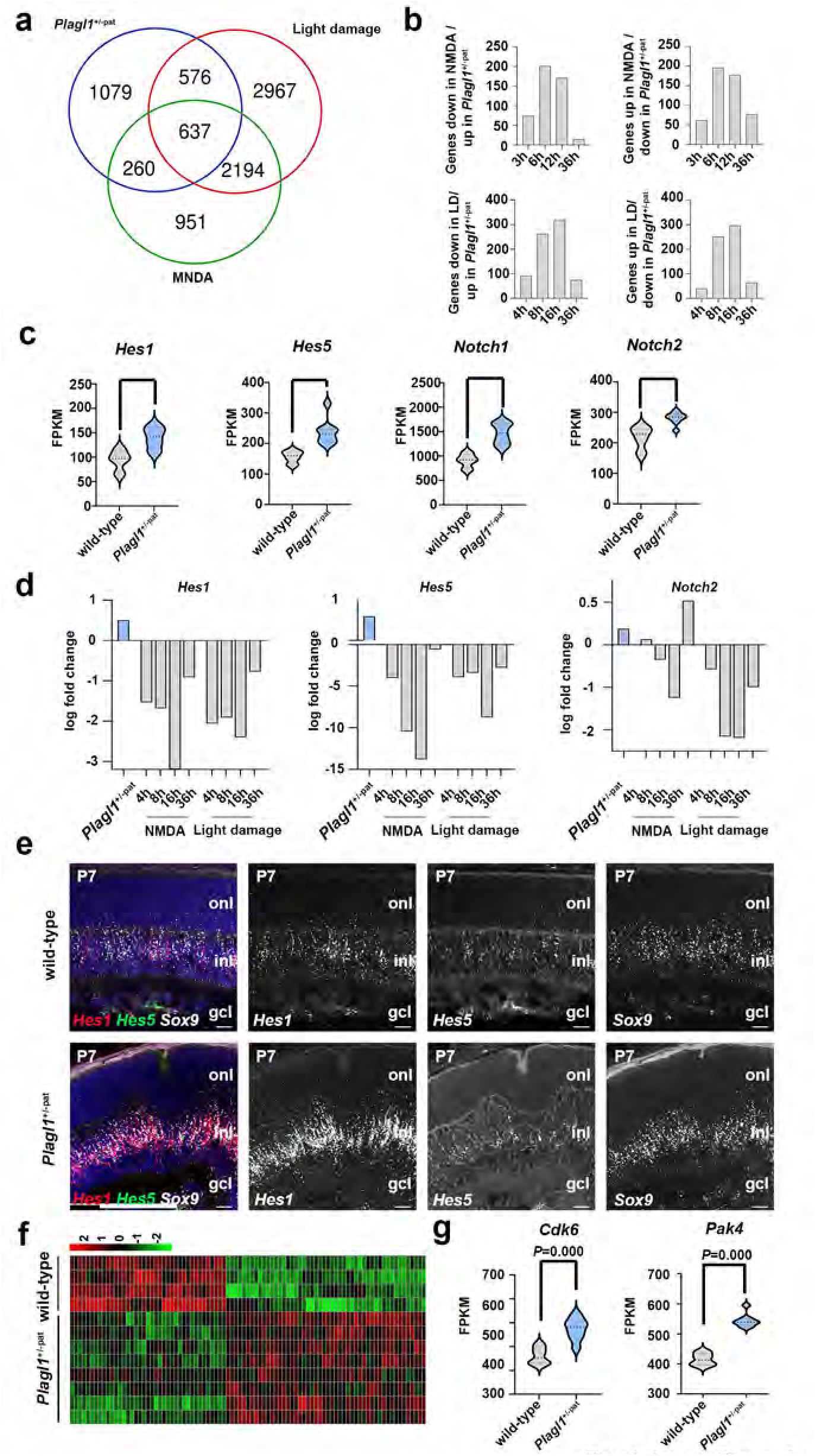
*Plagl1* inhibits Notch signalling and controls cell cycle genes in P7 Müller glia. **a** Venn diagram illustrating transcriptomic comparison of Müller glia after retinal treatment with NMDA/LD and *Plagl1*^+m/-pat^ retinas showing shared and specific DEGs in the 3 conditions. **b** Quantification and classification of shared DEGs in of NMDA/LD-activated Müller glia at different time points and *Plagl1*^m/-pat^ retinas that follow opposite transcriptional trajectories (up-in NMDA/LD Müller glia and down in *Plagl1*^+m/-pat^ and vice versa). **c** Violin plot showing increased expression of Notch pathway genes *Hes1, Hes5, Notch1*, and *Notch2* in P7 *Plagl1*^+m/-pat^ retinas. **d** Comparison of *Hes1, Hes5*, and *Notch2* transcript levels in *Plagl1*^+m/-pat^ retinas to NMDA/LD Müller glia at different time points post injury, comparing RNA-seq datasets. **e** RNAscope probe labelling of *Hes1, Hes5*, and *Sox9* on P7 retinal cross sections from wild-type and *Plagl1*^+m/-pat^ containing rosettes and showing *Hes1* and *Hes5* up-regulation is colocalized with *Sox9* staining. **f** Heatmap representing cell cycle genes differentially expressed in wild-type vs *Plagl1*^+m/-pat^retinas. **g** Violin plot showing up-regulation of cell cycle genes *Cdk6* and *Pak4* in P7 *Plagl1*^m/-pat^ retinas compared to wild-type.

To confirm that an increase in Notch signaling did occur in *Plagl1*^+/-pat^ Müller glia, we performed RNAscope on P7 control and *Plagl1*^+/-pat^ retinas. We hybridized labeled riboprobes to retinal sections to detect transcripts for *Hes1, Hes5* and the Müller glia marker *Sox9*. As expected, both *Hes5* and *Hes1* were expressed in the INL, co-localized with *Sox9*, and strikingly, their expression levels were increased in P7 *Plagl1*^+/-pat^ Müller glia (Fig. 6e). In addition, this up-regulation was confirmed by RT-qPCR confirmed in P7 *Plagl1*^+/-pat^ mutant retinas (Supplementary Fig. ^5d^). These data indicate that *Plagl1* loss of function results in Notch signalling pathway increase in P7 Müller glia.

### Upregulation of cell cycle genes in *Plagl1*^+/-pat^ retinas

In contrast to injured murine Müller glia that respond with limited or no proliferation, *Plagl1*^+/-pat^ Müller glia undergo robust proliferation at P7 (Fig. 4). We thus assessed the potential deregulation of cell cycle genes in *Plagl1*^+/-pat^ retinas by compiling a list of 616 genes involved in cell cycle regulation, including all genes in the GO category “GO0007049: cell cycle” and a functional classification of additional cell cycle genes based on GO categories and KEGG pathways using DAVID. Enriched categories were organized into clusters based on semantic similarity. Of the 616 compiled genes, 90 genes were deregulated (14.6%) in *Plagl1*^+/-pat^ retinas, including both down-regulated (50 genes) and upregulated (40 genes) genes, as depicted in a heatmap (Fig. 6f). Among the downregulated genes were known negative regulators of the cell cycle (*Cdkn2d*), while positive regulators were amongst the upregulated genes (*Cdk6, Cdk11b, Pard3, Pak4*) (Fig. 6f).

To identify potential regulators of Müller glia proliferation, we screened the 90 DEGs involved in cell cycle regulation for Müller glia specific genes. We identified 11 cell cycle genes specifically expressed in Müller glia and differentially expressed in *Plagl1*^+/-pat^. Amongst these genes *Rgs2, Arf6, Klh113* were down-regulated while *Cdk6, Src, Rcc2, Mad2l2, Pdcd6ip, Trp53bp2 and Pak4* were up-regulated in P7 *Plagl1*^+/-pat^ (Fig. 6g). 64 genes were expressed in Müller glia but not specific to, with 27 genes upregulated and 37 down regulated in *Plagl1*^+/-pat^ (Table 1). Finally, we analyzed the expression of these genes in NMDA/LD injured Müller glia and compared their transcriptional behaviour following injury to their expression in *Plagl1*^+/-pat^. Interestingly, *Pak4* was up-regulated in *Plagl1*^+/-pat^ and not differentially expressed in NMDA/LD injured Müller glia. In contrast, *Numa1, Cep131* were up-regulated in *Plagl1*^+/-pat^ and down-regulated in NMDA/LD injured Müller glia, while *Rps3, Rab11a, Zwint, Uba3* and *Lig4* were down-regulated in *Plagl1*^+/-pat^ and up-regulated in NMDA/LD injured Müller glia (Supplementary Fig. ^6a-b^).

Taken together these data support the notion that the loss of *Plagl1* function results in a robust proliferation of Müller glia at P7 in the retina. Furthermore, our transcriptional analysis suggests *Plagl1* acts upstream genes and signalling pathways that control Müller glia proliferation especially Notch signalling pathway genes, as well as *Lhx2*, and Yap genes *Tead1,2* and a network of cell cycle genes.

## DISCUSSION

In the retina, *Plagl1* is expressed in retinal progenitor cells, where it plays a role as a negative regulator of cell number control, ensuring that appropriate numbers of rod photoreceptors and amacrine cells are generated ^24^. In addition, *Plagl1* is expressed in subsets of mature retinal cells, including adult Müller glia ^62^, where its functions have not been studied. Here we found that *Plagl1* is expressed in a dynamic fashion in Müller glia post-injury, with a rapid and transient upregulation that is followed by a more long-term decline. To study *Plagl1* function, we examined *Plagl1*^+/-pat^ animals, which we confirmed to be null mutants, revealing that they develop defects in retinal architecture and visual function beginning at perinatal stages, which correlate with a reactive gliotic phenotype of Müller glia. Furthermore, we found that *Plagl1*^+/-pat^ Müller glia proliferate in an injury-independent fashion, and give rise to new inner retinal neurons and photoreceptors post-injury. Based on transcriptomic analyses, we found that *Plagl1* is a negative regulator of several pro-proliferative and pro-glial pathways in the retina, including Notch signaling. The loss of *Plagl1* thus initiates the three critical steps in regeneration in murine Müller glia, which include reactive gliosis, re-entry into the cell cycle, and neurogenesis.

In addition to the shared structural abnormalities between *Plagl1*^+/-pat^ and other retinal degenerative models, we observed several overlapping transcriptomic changes with *Ndp, rd10*, and *rds* mutants, as well as some overlap with the NMDA injury model. These findings are in keeping with the central role that Müller glia dysfunction plays in retinal degeneration, including in human disease, including in macular telangiectasia type 2 (MacTel-2) ^63^. We attribute the *Plagl1*^+/-pat^ degeneration-like phenotype to a requirement for *Plagl1* in Müller glia because the overlapping phenotypes observed after Müller glia are ablated, including retinal rosettes, vascular defects, and abnormal ERG recordings ^38^.

The role of *Plagl1* in sustaining quiescence is consistent with its role as a tumor suppressor gene. In humans, *PLAGL1* localizes to a chromosomal region (6q24-25) linked to growild-typeh inhibition, its ability to induce cell cycle arrest and apoptosis when misexpressed in epithelial cell lines ^51,53^, and its inactivation in breast ^53^, pituitary ^64^, head and neck ^65^, and ovarian carcinomas ^66,67^. However, our study provides the first evidence of an *in vivo* role for *Plagl1* in maintaining non-malignant somatic cells in quiescence. *Plagl1* null mutant retinas also resemble several other Müller glia-specific conditional KOs (cKOs), such as *Dicer* cKOs, which also have retinal dysmorphology and defects in retinal function, although hey do not display ectopic proliferation^68^. Transcription factors of the Nfi family can also promote a glial identity and cell cycle exit when misexpressed in late RPCs ^60^. Interestingly, misexpression of murine *Plagl1* in Xenopus ^23^ and mouse ^24^ also promotes a Müller glial cell fate, but only in mouse is PLagl1 necessary and sufficient to promote cell cycle exit. Taken together with this study, we conclude that Plagl1 plays a role not only in the development of Müller glia, but also in maintaining the quiescence of these cells at postnatal stages.

However, ectopic proliferation of *Plagl1*^+/-pat^ Müller glia was transient, only observed at P7 and not later, and even at P7, not all Müller glia had re-entered the cell cycle, even though most of these cells express *Plagl1*. This finding is similar to the transient proliferative response of mammalian Müller glia to injury ^69-71^, and suggests that there may be distinct populations of Müller glia that respond to injury. In addition, there is growing support that there are active inhibitory processes that have to be overcome for Müller glia to re-enter the cell cycle. One example is *Cdkn1c*, which plays a critical role in preventing cell cycle re-entry of Müller glia. However, *Cdkn1c* was not downregulated in *Plagl1* mutant retinas, although *Cdkn2d*, another cell cycle inhibitor, was reduced in expression, and multiple positive cell cycle regulators were also elevated (Fig. 6f).

Another conundrum is the increased expression of *Nfib/c/x* in *Plagl1* mutant retinas, which promote a glial identity in the retina ^60^, but are also required to promote quiescence, although injury response can induce ectopic proliferation in these mutants ^7^. In contrast, in this study, we showed that Müller glia proliferate ectopically in *Plagl1*^+/-pat^ retinas even without injury. Similarly, *Lhx2* is required to prevent reactive gliosis in an injury-independent fashion^72^, but we found that *Lhx2* is upregulated in *Plagl1* mutants. One possibility is that *Plagl1* acts downstream of *Nfi* and *Lhx2* transcription factors, such that its loss has a dominant action in maintaining Müller glia quiescence.

*Plagl1* is a maternally imprinted gene that is expressed in a parent-of-origin-specific fashion (i.e., paternal allele only) due to DNA methylation of the maternal allele, a modification that first appears in the germ cells^73^. Imprinting affects < 1% of all genes in placental animals, including mammals^74,75^. Imprinted genes are heterogeneous in function (e.g., transcription factors, signaling molecules, cell cycle regulators), but recent studies suggest they may share a common function in the maintenance of cellular quiescence^27^. A meta-analyses of 85 known IGs, revealed that these genes are co-regulated; *Plagl1* is part of an IGN of 13 imprinted genes (*Meg3, Ndn, Grb10, Dlk1, Igf2, Cdkn1c, Zac1, Peg3, Mest, Nnat, Asb4, H19, Ppp1r9a*) that are tightly co-regulated^27,31^. Strikingly, many of these IGN genes are negative regulators of cell growild-typeh (Igf2 a notable exception) whether the imprint is maternal or paternal^27,31^. Of these, *Dlk1*^29^ and *Cdkn1c*^30^ maintain adult neural stem cell (NSC) quiescence in the forebrain. Notably, two IGN genes become biallelically expressed in adult somatic cells; *Dlk1* in adult NSCs^29^; *Igf2* in meninges, choroid plexus and vasculature^76^. In contrast, our analysis of transcript reads revealed that *Plagl1* remains mono-allelically expressed in the postnatal retina.

Expression of several of these IGN genes is also associated with cell cycle withdrawal/differentiation *in vitro*^27^. However, our work demonstrating that *Plagl1* maintains retinal Müller glia in quiescence is the first to establish this relationship *in vivo*. Notably, *Plagl1* mutant MEFs do not proliferate ectopically^49^, suggesting that *Plagl1* sustains quiescence only in a subset of somatic cells *in vivo*, and one intriguing possibility is that this function is restricted to those cells with stem cell potential. Consistent with this idea, two members of the IGN, *Dlk1*^29^ and *Cdkn1c*^30^, maintain adult neural stem cell quiescence in the forebrain. Interestingly, of the imprinted genes in the *Plagl1* IGN, only *Dlk1* was differentially expressed in *Plagl1*^+/-pat^ retinas, with an increase in transcript levels. *Dlk1* is a negative regulator of Notch signaling ^77^, but as multiple downstream effectors of Notch are increased in *Plagl1*^+/-pat^ retinas, it does not appear that *Dlk1* has a functional role in inhibiting Notch signaling in these animals. Notably, *Dlk1* (also known as *Pref1*) can activate ERK signaling in adipocytes ^78^, and as ERK signaling is upregulated in *Plagl1*^+/-pat^ Müller glia, *Dlk1* may contribute to this process. ERK signaling is activated downstream of receptor tyrosine kinases (RTKs) in response to growth factor signaling (e.g., EGF, IGF, FGF2), and activation of this pathway is a universal feature of reactive gliosis in all species^6^.

In conclusion, we have identified *Plagl1* as a critical regulator of mammalian Müller glia quiescence, one of the few genes that when mutated initiates a proliferative glial response. There are clear implications for regenerative medicine, in which the goal is to initiate the endogenous Müller glial repair response that occurs in fish and frogs.

## METHODS

### Animals

Animal procedures were approved by the University of Calgary Animal Care Committee (AC11-0053) and later by the Sunnybrook Research Institute (16-606) in compliance with the Guidelines of the Canadian Council of Animal Care and conformed to the ARVO statement for the Use of Animals in Ophthalmic and Vision Research. To stage embryos, we considered the morning of the vaginal plug as embryonic day (E) 0.5. *Plagl1*^+/-pat^ animals were generated by crossing *Plagl1*^+/-mat^ males (obtained from Laurent Journot ^31^) with C57/Bl6 wild-type females (#000664, Jackson Laboratory, ME, USA). Genotyping was performed with the following primers: *Plagl1* mutant allele: forward (5’-ATG GCT TCT GAG GCG GAA AG-3’) and reverse (5’-AAA GGC TCC AAA GGC TCC AAG G-3’); *Plagl1* wild-type allele: forward (5’-TCG TCA CAC CAA GAA GAC CCA C-3’) and reverse (5’-AAA GGC TCC AAA GGC TCC AAG G-3’); CD1 mice (Charles River Laboratories, Senneville, QC, Canada) were used for explant experiments.

### Electroretinogram (ERG)

Animals were dark-adapted for one hour prior to ERG recordings. ERGs were recorded under dim red light. Stimulation and acquisition were performed with an Espion E^2^ system (Diagnosys LLC) (flash duration 10 μs, bandpass filtering 0.3Hz-3Khz), as described ^79,80^. Briefly, scotopic intensity responses were measured using increasing light flash pulses of increasing strength (−5.22 to 2.86 log cds/m^2^) and photopic intensity responses (30 cd/m^2^ background light) used increasing flash strengths from −1.6 to 2.9 log cds/m^2^.

### Optical coherence tomography (OCT)

Mice were anesthetized with 2% isoflurane general anaesthesia after their pupils were dilated using tropicaminde topical drops (Mydriacyl 1%; Alcon Canada Inc). Eyes were imaged using the Spectralis optical coherence tomography (OCT) and confocal scanning laser ophthalmoloscopy (cSLO) system mounted with an additional 25-diopter lens (Heidelberg Engineering, Germany). Normal saline was applied to the mice corneas every 2 minutes to keep it hydrated. OCT was performed on a 30°× 20° area of P21 mouse retinas. A total of 31 scans were acquired with an average of 20 frames per scan. Infrared cSLO scans (IR-cSLO) were acquired (**λ** = 815 nm) with an average of 30 frames per scan to assess the retinal fundus of Zac1 mutant and controls. Retinal scans were exported from the HEYEX software (Heidelberg Engineering, Germany) as tiff images and processed using ImageJ software (NIH, USA).

### Tissue processing

Retinal tissues were drop fixed in 4% paraformaldehyde (PFA)/1X phosphate buffered saline (PBS) at 4°C overnight for dissected retinas or eyes or 3 hours for retinal explants. Fixed tissues were then rinsed 3 × 10 min in 1X PBS and cryopreserved in 20% sucrose/1X PBS overnight at 4°C. For cryosectioning, tissues were embedded in O.C.T™ (Tissue-Tek®, Sakura Finetek U.S.A. Inc., Torrance, CA). Cryosections were cut at 10-12 microns on a Leica CM3050s cryostat (Leica Biosystems, Buffalo Grove, IL, USA) and collected on Fisherbrand™ Superfrost™ Plus slides (Thermo Fisher Scientific, Markham, ON). Collected slides were either processed directly or stored at −20°C in a tape-sealed slide box.

### Immunohistochemistry

Cryosections were blocked for 1 hr at room temperature in blocking solution: 10% normal goat serum in PBST (1X PBS/0.1% Triton X-100). Primary antibodies were diluted in blocking solution and incubated at 4ºC overnight. The following primary antibodies were used: rabbit anti-Arr3 (1/500, Millipore #AB15282), rat anti-BrdU (1/500, Serotec #OBT0030S), mouse anti-Brn3a (1/500, Chemicon #5945), mouse anti-calbindin (1/1000, Sigma #9848), rabbit anti-calretinin (1/2000, Swant #76699/4), rat anti-CD11b (M1/70, 1/500, Abcam #ab8878), mouse anti-Rlbp1 (1/500, Abcam #15051), rabbit anti-GFAP (Sigma #G9269), rabbit anti-Glul (1/500, Abcam #73593), rabbit anti-Iba1 (1/500, Cat#019-19741), rabbit anti-M-opsin (1/250, Millipore #5405), rabbit anti-Pax6 (1/500, Convance #PRB-278P), Rabbit anti-pERK (Map kinase p44/42 phospho-Thr202/Tyr204) (1/500, Cell signaling #4370), mouse anti-rhodopsin (1/500, Chemicon #MAB5356), rabbit anti-S-opsin (1/250, Millipore #5407), rabbit anti-Sox9 (1/500, Millipore #AB5535), mouse anti-syntaxin (1/500, Sigma #S 0664), mouse anti-vimentin (1/500, Sigma #V5255), or anti-ZO-1 (1/100, ThermoFisher #33-9100), Slides were washed 3 × 10 min in PBST and then incubated in secondary antibodies conjugated to Alexa-568, Alexa-488 or Alexa-647 diluted at 1/500 in PBST for 1 hr at room-temperature, followed by 3 × 10 min washes in PBST.

### Isolectin staining

Eyes were dissected and retinas were flatmounted or processed for sectioning as described. Wholemount retinas were fixed in 4% PFA/1X PBS for 2 hours. Flatmounted retinas or cyrosections were fixed in ice cold 70% ethanol for 30 min. Fixed tissue was then incubated in Isolectin B4 (1/250, Sigma #L2140) in PBST overnight at 4°C, washed 3 × 10 min in PBST. Wholemount retinas were mounted on microscope slides and mounted tissues or sections were coverslipped in Aquapolymount.

### BrdU labeling

To label S-phase cells, animals were injected intraperitoneally with 100 g/g body weight BrdU (Sigma, Oakville ON) either 30 min before sacrifice for proliferation assays or at P7 for birthdating assay. Tissues were processed for anti-BrdU staining as described above except for the addition of a pre-treatment step with 2N HCl for 15 min at 37°C prior to blocking.

### Retinal explants

Retinas were dissected and the RPE removed from P0 eyes in ice-cold PBS. Retinas were flattened on 0.25 μm Nucleopore membranes (Whatman # 110409) in 6 well plates and cultured for 14 days in vitro (DIV) at 37°C, 5% CO_2_ in retinal explant medium. (50% Dulbecco’s Modified Eagle Medium (DMEM, Wisent; #319-005-CL), 25% Hanks’ Balanced Salt Solution (HBSS, Gibco; #24020-117), 25% heat inactivated horse serum (ref), 200 μM L-Glutamine (Wisent; #609-065-EL), 0.6 mM HEPES (Wisent; # 330-050-EL), 1% Pen/Strep (Wisent; # 450-201-EL). For lineage tracing, 4-hydroxytamoxifen (4-OHT) (Sigma-Aldrich; #H6278) was added to the retinal explant media after 7 DIV at 5 mM final concentration. Half of the media was removed and replaced with fresh media every day with a 2X concentration of 4-OHT. Explants were harvested after 14 DIV.

### RNA extraction

Total RNA was extracted from dissected retinas using Trizol® RNA Isolation Reagent (Thermo Fisher Scientific; #15596-026), following the manufacturer’s instructions except the overnight incubation step was reduced to 30 min at −80°C. RNA quality was assessed using an Agilent 2100 Bioanalyzer (Agilent Technologies, Palo Alto, CA, USA).

### RNA sequencing

For RNA-Seq 8 *Plagl1*^+/-pat^ and 4 wild-type littermates were analyzed. 500 ng RNA per sample was sequenced on Illumina NextSeq500 Platform. 75 base single-end sequence reads were generated on NEBNext Ultra II Directional RNA Library Prep Kit for Illumina. Basecalling and demultiplexing was done using IIllumina CASAVA 1.9 pipeline. Reads were mapped to mouse genome (Ensembl, GRCm38) using hisat2 version 2.1.0 with a mapping rate close or over 98% in each sample.

### Differential expression and clustering analysis

Differentially expressed genes were detected using DESeq2 1.24.0 Bioconductor package https://bioconductor.org/packages/release/bioc/html/ DESeq2.html (Love et al., 2014, p. 2) as described in package vignette. Genes with adjusted p-values (Benjamini-Hochberg adjustment for multiple comparisons) (Benjamini and Hochberg, 1995) less then 0.05 (5% chance of gene being a false positive) and over 1.5 fold change in either direction were selected as differentially expressed.

### Gene Ontology analysis

Over-represented gene ontology (GO) terms and Kyoto Encyclopedia of Genes and Genomes (KEGG) (Kanehisa and Goto, 2000) pathways were detected using clusterProfiler 3.12.0 Bioconductor package (Yu et al., 2012). The analysis was done separately was on the lists of up- and down-regulated genes.

### ATAC-seq

ATAC-seq was performed using 75,000 cells as input material for the library preparation with Active motif kit (Catalog No. 53150) following the manufacturer’s instructions. 75 bp paired-end sequencing was performed on Illumina NEXTSEQ500. Reads were mapped to the reference genome (Ensembl, GRCm38) using bowild-typeie2 short read aligner version 2.3.5.1. Areas of open chromatin were predicted using Macs2 v.2.2.7.1 based on alignment files filtered of mitochondrial reads. Differential enrichment was detected using DiffBind Bioconductor package. Data was normalized using Trimmed Mean of Medians (TMM) normalization native to EdgeR Bioconductor package. In total, differential binding tests were applied to 56771 peaks. The peak had to be present in at least 2 samples within experimental group to be included into the analysis. False discovery rate (FDR) below 0.05 was a threshold to consider the peak differentially bound. No peaks were below 0.05 FDR threshold. DiffBind results were annotated with nearest promoter, gene symbol, entrez_id and gene description using ChIPpeakAnno Bioconductor package. Term enrichment analysis was conducted using 2 different approaches: 1) over-representation analysis (ORA) conducted using geneSCF software and 2) Gene set enrichment analysis (GSEA). Genes with TSS located within 5000 bp of DARs were used in ORA. The input for GSEA was the list of all genes within 5000 bp from the nearest TSS sorted by the Fold Change. The GSEA enrichment analysis was conducted using ClusterProfiler v.3.18.1 Bioconductor package.

### ATAC-seq and Zac1 RNA-seq integration

ATAC-seq and RNA-seq were integrated by overlapping the lists of significantly changed genes and DARs. Genes or DARS with adjusted p-values below 0.05 were considered significant. The list of DARs was limited to 311 peaks located within 5000 bp from the nearest TSS.

### RT2 Real-time Quantitative Reverse Transcription-PCR

For complimentary DNA (cDNA) synthesis, 0.5 μg of total RNA was converted using an RT^2^ first strand kit (Qiagen, #330401) following the manufacturer’s instructions. qPCR reactions were performed using a CFX384 cycler (Biorad Laboratories, Canada) using a RT^2^ SYBR® Green PCR Master Mix (Qiagen; #330500) following the manufacturer’s instructions. The following RT^2^ qPCR primers were used: *Sox9* (PPM05134D), *Lhx2* (PPM31533A), *Hes1* (PPM5647A), *Hes5* (PPM31391A) and three reference genes for normalization: *Gapdh* (PPM02946E), *B2m* (PPM03562A), and *Hrpt* (PPM03559F). The *ΔΔ*Ct method was used to determine relative gene using the Biorad CFX manager software.

### RNAscope

Double fluorescent *in situ* hybridization were performed using a RNAscope^®^ Multiplex Fluorescent Detection Kit v2 (ACD; #323110) according to the manufacturer’s directions. ACD probes used included: Mm-*Sox9* (C2: 401051-C2), Mm-*Hes1* (#417701), Mm-*Hes5* (400991-C2), and Mm-*Plagl1* (C1: 462941). Opal™ 570 (Akoya; #FP1488001KT; 1:1500) was applied for channel 1 and Opal™ 520 (Akoya; #FP1487001KT; 1:1500) was applied for channel 2. Retinal sections were counterstained with DAPI and mounted in Aquapolymount as described.

### Western blotting

Retinas and retinal explants were collected from adult and postnatal pups at the indicated stages, lysed in RIPA buffer with protease (1x protease inhibitor complete, 1 mM PMSF) and phosphatase (50 mM NaF, 1 mM NaOV) inhibitors, and 10μg of lysate was run on 10% SDS-PAGE gels for Western blot analysis as described previously (Ma et al., 2007). Primary antibodies included Gfap (Sigma, #G9269, 1/10.000), Vimentin (Sigma, #V5255, 1/10.000), pERK, Glutamin Synthetase (Abcam, #ab73593, 1/10.000) and Actin (Abcam; #ab8227, 1/10.000). Densitometries were calculated using ImageJ. The average values of normalized expression levels were plotted.

### Image acquisition and processing

Images were captured on a Leica DMRXA2 optical microscope (Leica Microsystems Canada Inc., Richmond Hill, Ontario, Canada) using LasX software. Confocal images were acquired using a Leica SP8 spectral confocal microscope (Leica Microsystems CMS GmbH). The images were prepared using Adobe Photoshop CC 2017 (Adobe Systems Inc., San Jose, CA, USA). For *Plagl1*^+/-pat^ retinal transverse sections, all areas and sections containing rosettes and/or ectopia were imaged, the equivalent areas/ sections number were imaged for the wild-type littermate and used as controls.

### Statistical analysis

Cell counts were performed on a minimum of three retinas for each genotype, and a minimum of three sections from each eye. The n numbers and statistical tests used for each count were indicated in the figure legends. Graphs and statistics were generated using GraphPad Prism Software 8 (GraphPad Inc., La Jolla, CA). All data expressed as mean value ± standard error of the mean (SEM). In all experiments, a p value < 0.05, **p < 0.01, ***p < 0.005, and ****p < 0.001.

## ACKNOWLEDGEMENTS

CS holds the Dixon Family Chair in Ophthalmology Research at Sunnybrook Research Institute. This project is currently supported by funding to CS from Natural Sciences and Engineering Research Council of Canada Discovery Grant-RGPIN-2017-03649 and Canadian Institutes of Health Research (CIHR) Operating Grant MOP-142338. YT was supported by a U of Calgary Eyes High Fellowship, MH by an Alberta Innovates Health Solution (AIHS) summer studentship, LA and NT by a CIHR/Alberta Children’s Hospital Research Institute training grant, and RD by a CIHR Canada Hope Fellowship.

## AUTHOR CONTRIBUTIONS

YT: conceptualization, data curation, formal analysis, investigation, methodology, visualization, validation, writing – original draft, writing – review and editing

LAD: formal analysis, investigation, methodology, visualization, validation, writing – review and editing

YI: data curation, formal analysis, software

EvO: data curation, formal analysis, software

JH: formal analysis, investigation, methodology, visualization, validation, writing – review and editing

NT: formal analysis, investigation, methodology, visualization, validation

LA: data curation, formal analysis, investigation

JZ: data curation, formal analysis, investigation

MH: data curation, formal analysis, investigation

RD: data curation, formal analysis, investigation

LJ: resources, writing – review and editing

YS: resources, data curation, formal analysis, investigation, writing – review and editing

IK: resources, supervision, software, writing – review and editing

IA: funding acquisition, resources, supervision, validation, writing – review and editing

JB: resources, supervision, validation, writing – review and editing

CS: funding acquisition, conceptualization, project administration, resources, supervision, validation, writing – original draft; writing – review and editing

## FIGURE LEGENDS

**Supplementary Figure 1.**
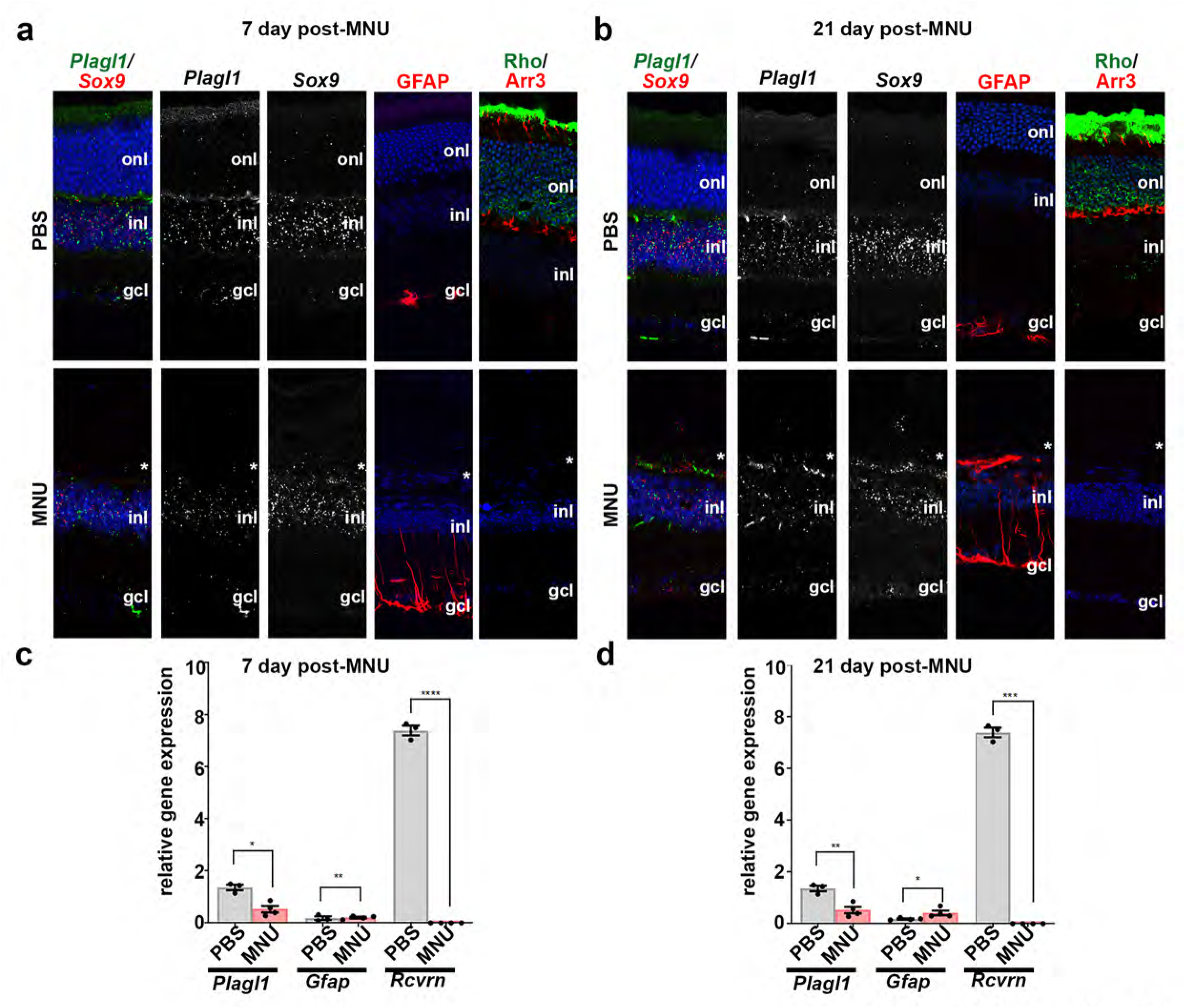
*Plagl1* expression is down-regulated in Müller glia following injury. **a**,**b** RNAscope staining for *Plagl1, Sox9*, and immunofluorescence for GFAP, Rho and Arrestin3 in P7 wild-type retinal section of mice treated with PBS and MNU 7(a) - and 21(b) -days post injection. **c**,**d** Quantification of *Plagl1, GFAP* and *Recovrin* expression by qRT-PCR in wild-type retinas treated with PBS and MNU and collected after 7-days **(c)** and 21-days **(d)** post injection. N=3 for all time points. Statistics: mean ± S.E.M, Student t-test. significant differences: *p<0.05, **p<0.01, ***p <0.0001. gcl, ganglion cell layer; inl, inner nuclear layer; onl, outer nuclear layer. Scale bars: 50 μm.

**Supplementary Figure 2.**
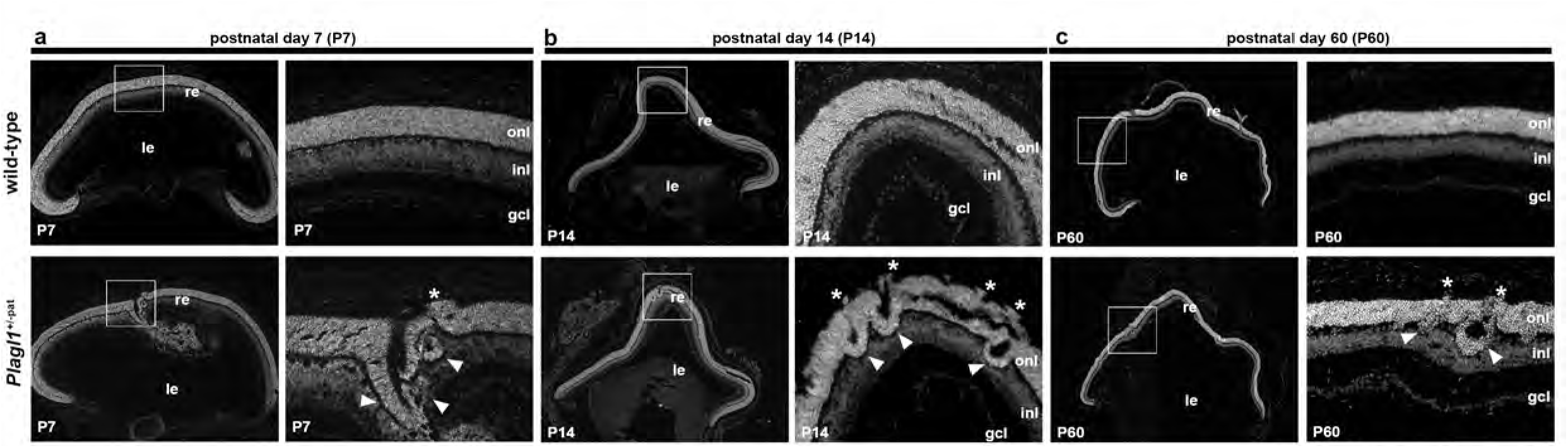
*Plagl1* is required to sustain retinal integrity. **a-c** DAPI nuclear staining of P7 (a), P14 (b), and P60 (c) wild-type and *Plagl1*^+/-pat^ retina sections, showing retinal ectopias (asterisks) and rosettes (arrowheads). Right side panels are high magnifications of boxed areas. gcl, ganglion cell layer; inl, inner nuclear layer; le, lens; onl, outer nuclear layer; re, retina.

**Supplementary Figure 3.**
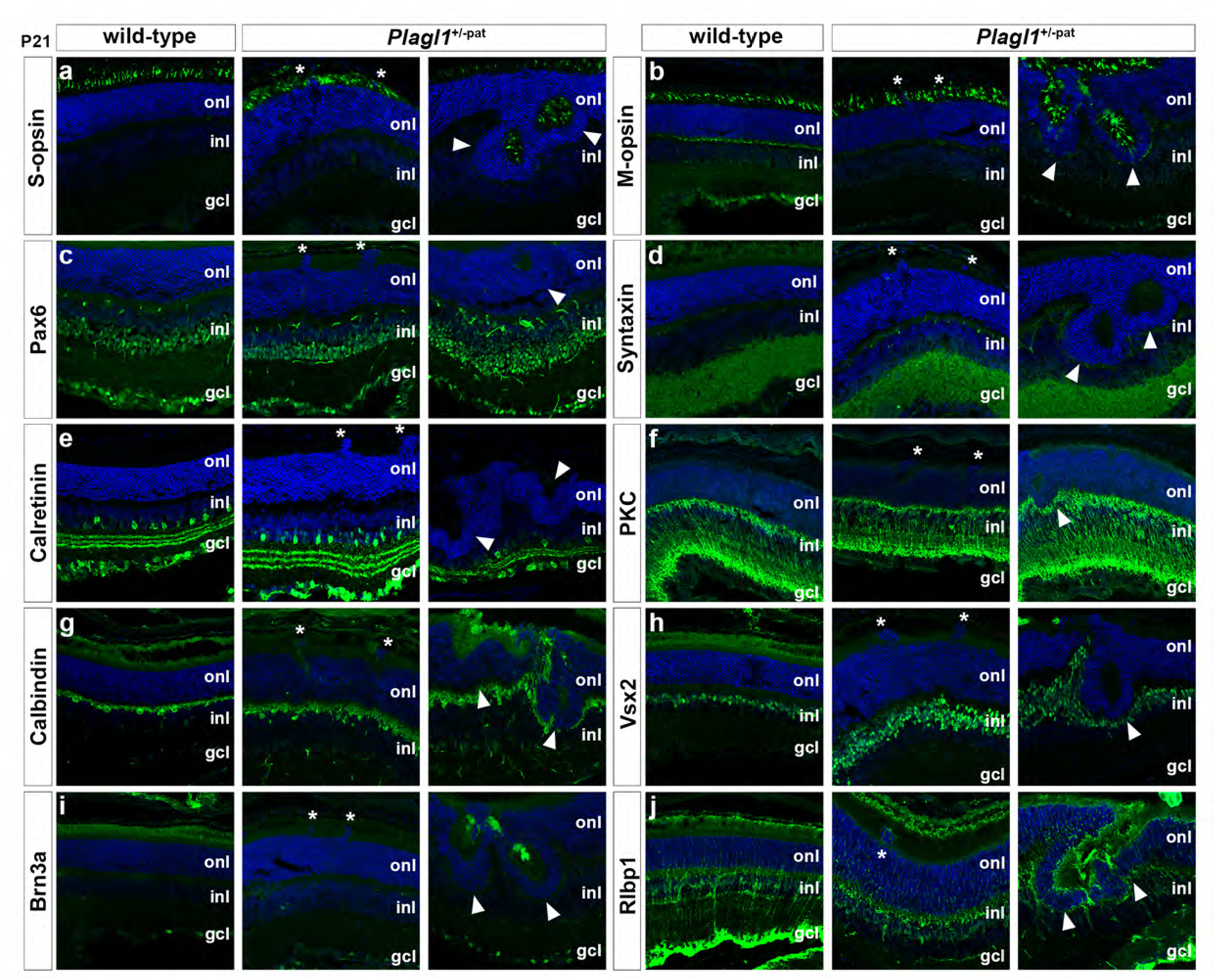
All retinal cell types are present in *Plagl1*^+m/-p^ retinas. **a-j** Immunofluorescence for retinal cell specific markers S-opsin (cones), Pax6 (amacrine and ganglion cells), Calretinin (amacrine cells), Calbindin (horizontal cells), Brn3a (ganglion cells), M-opsin (cones), Syntaxin (amacrine cells), PKC and Vsx2 (bipolar cells), and Rlbp1 (Müller glia) with DAPI nuclear counter-staining of P21 wild-type and *Plagl1*^+m/-pat^ retinal sections, showing ectopia (Asterisks) and rosettes (Arrow heads) contain all retinal cell types. At least 3 *Plagl1*^+m/-pat^ mice of 2 different litters and their wild-type littermates were analyzed. gcl, ganglion cell layer; inl, inner nuclear layer; onl, outer nuclear layer.

**Supplementary Figure 4.**
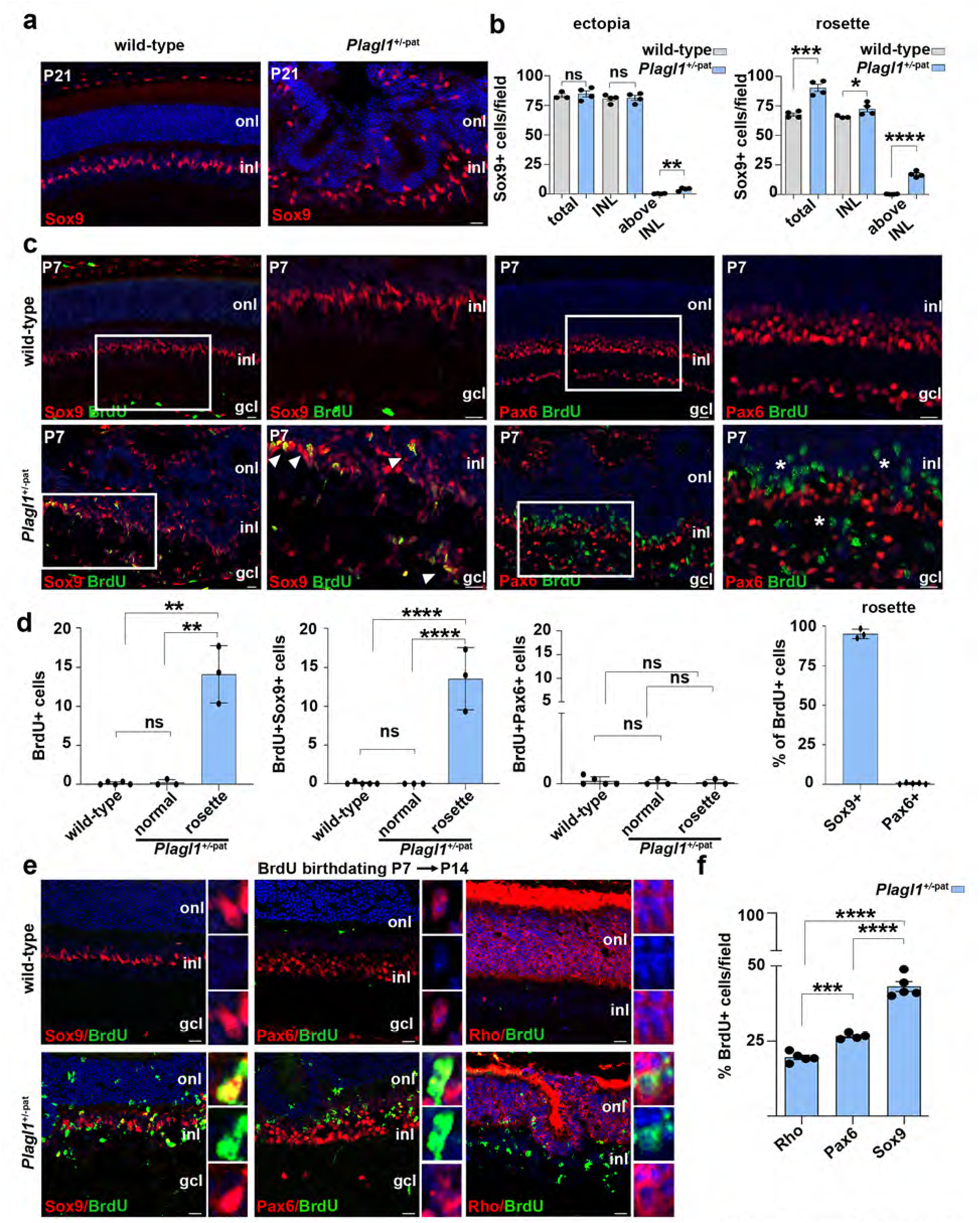
GO term and KEGG pathways analysis of differentially expressed genes in the *Plagl1*^+m/-p^ transcriptome. **a** Gene ontology and KEGG analysis identifying molecular functions, biological processes, cellular compartments and pathways deregulated in P7 *Plagl1*^+m/-p^ retinas. **b** Gene ontology analysis showing the top cellular compartments and molecular functions comparing *Plagl1*^+m/-pat^ retinal transcriptome to *Plagl1* CHIP-seq, *Plagl1* GOF in the cortex, *Plagl1*^+m/-pat^ MEF transcriptome, and *Plagl1* GOF in N2A cell data sets. **c** Transcriptomic comparison of *Plagl1*^+m/-pat^ DEGs vs *Norrin KO* DEGs **d** Transcriptomic comparison of *Plagl1*^+m/-pat^ DEGs vs *Rds KO* DEGs **e** Transcriptomic comparison of *Plagl1*^+m/-pat^ DEGs vs *Rd10 KO* DEGs

**Supplementary Figure 5.**
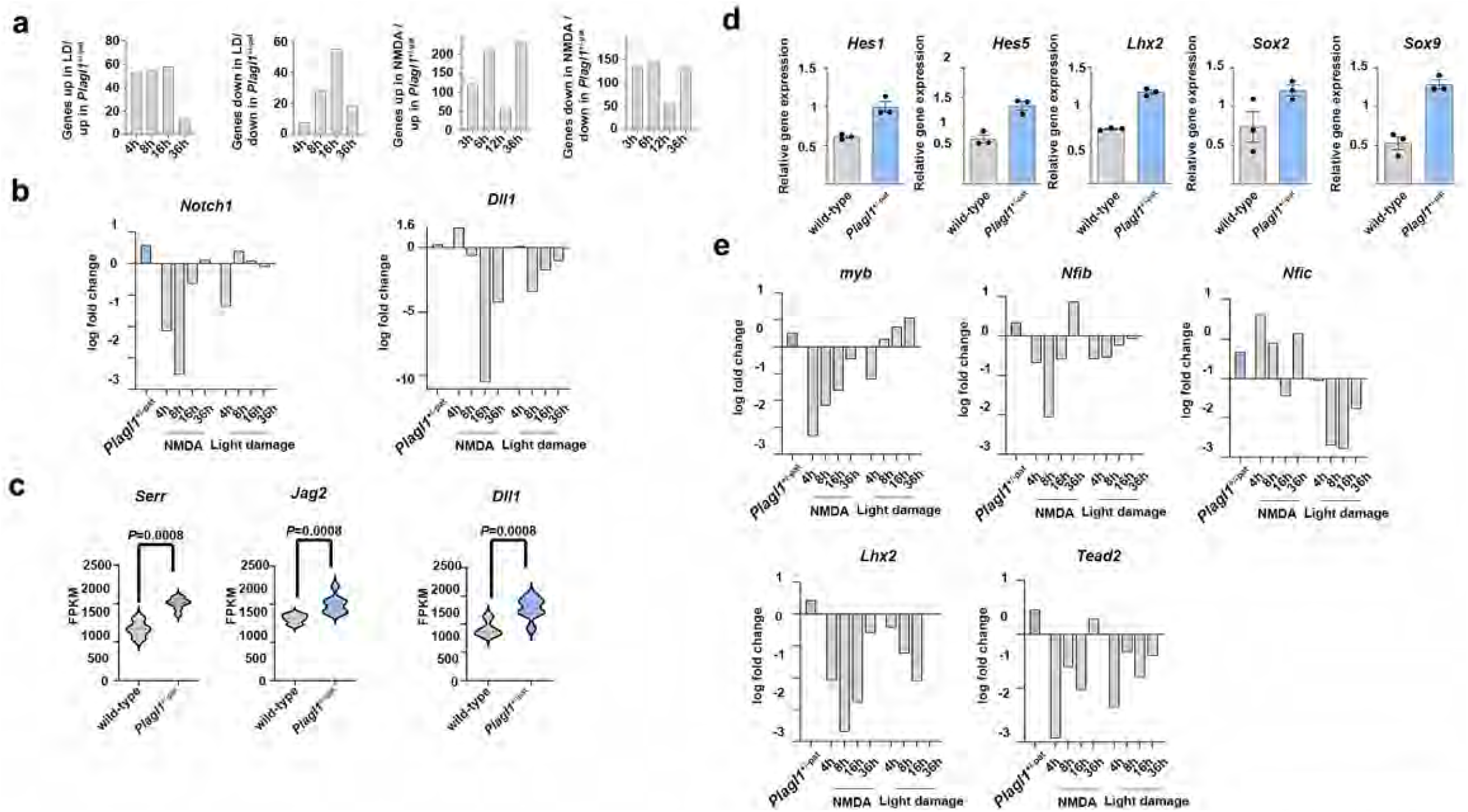
Cell cycle gene GO terms and KEGG pathways analysis in *Plagl1*^+m/-p^ transcriptome. **a** Transcriptomic comparison of NMDA/LD Müller glia and *Plagl1*^+m/-pat^ showing shared DEGs that follow the same trajectory, up-regulated, or down-regulated in the 3 conditions at different time points. **b** Notch signalling pathway genes *Notch1* and *Dll1* expression in Müller glia from NMDA/LD treated retinas and *Plagl1*^+m/-pat^ RNAseq a data set.s **c** Expression of Notch genes *Serr, Dll1* and *Jag2* in P7 *Plagl1*^+m/-pat^ RNAseq data set. **d** RT-qPCR expression analysis of Notch genes *Hes1, Hes5* and the gliogenic factors *Lhx2, Sox2, Sox9* in P7 *Plagl1*^+m/-pat^ compared to wild-type. **e** Transcriptomic comparison of Müller glia from NMDA/LD treated retinas and *Plagl1*^m/-pat^ retinas showing expression of genes involved in Müller glia reprograming *Myb, Nfib, Nfic, Lhx2*, and *Tead2*.

**Supplementary Figure 6.**
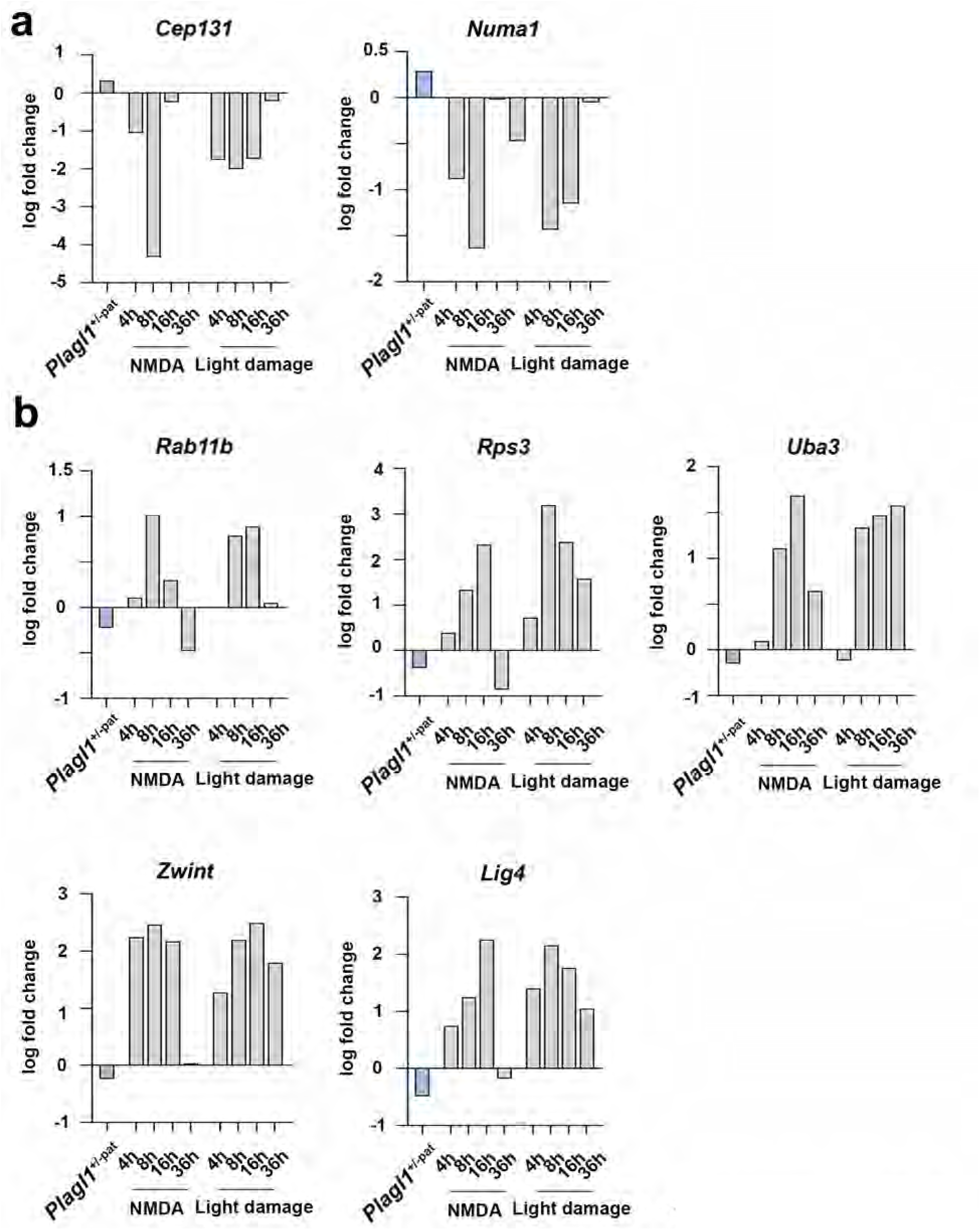
Cell cycle genes deregulated in P7 *Plagl1*^+m/-p^ retinas. **a** Transcriptomic comparison of NMDA/LD Müller glia and *Plagl1*^+m/-pat^ showing cell cycle genes whose expressions are up-regulated in *Plagl1*^+m/-pat^ while down-regulated in NMDA/LD Müller glia; *Cep131, Numa1*. **b** Transcriptomic comparison of NMDA/LD Müller glia and *Plagl1*^+m/-pat^ showing cell cycle genes whose expressions are down-regulated in *Plagl1*^+m/-pat^ and are initially up-regulated in NMDA/LD Müller glia; *Lig4, Rab11b, Rps3, Uba3, Zwint*

